# clusIBD: Robust Detection of Identity-by-descent (IBD) Segments Using Unphased Genetic Data from Poor-quality DNA Samples

**DOI:** 10.1101/2025.03.13.643019

**Authors:** Ran Li, Yu Zang, Zhentang Liu, Jingyi Yang, Nana Wang, Jiajun Liu, Enlin Wu, Riga Wu, Hongyu Sun

## Abstract

Inferring relatedness through the detection of identity-by-descent (IBD) segments is widely used in many fields. However, existing methods are often powerless for poor-quality DNA samples. Here, we propose a method, clusIBD, which can robustly detect IBD segments using unphased genetic data with high genotype errors. We evaluated and compared the performance of clusIBD with IBIS, TRUFFLE, and IBDseq using simulated data, artificial poor-quality materials and ancient DNA samples. The results showed that clusIBD outperformed these state-of-the-art existing tools and may be a promising tool for kinship inference in fields such as ancient DNA analysis and criminal investigation (freely available at https://github.com/Ryan620/clusIBD/).

## Background

Inferring relatedness from genomic data is an essential component in many fields, such as genetic association studies, population genetics, forensics, archaeology and ecology [1–6]. Specifically, inference of relatedness between samples is helpful to avoid spurious signals in genetic association studies because kinship among the cases or controls that is not known to the investigator may inflate the false positive rate [4]. In population genetic analyses, kinship needs to be taken into account, and sometimes relatives should be removed before other analyses are performed, such as population structure, human evolutionary history, times of population splitting, migration rates, and patterns of mating among individuals [7,8]. In forensic genetics, familial searching and investigative genetic genealogy can aid in finding criminals and identifying unknown deceased persons through DNA of their relatives [9–11], which extends the investigative lead value of current forensic DNA databases and Direct-to-Consumer (DTC) databases [12]. Kinship inference has also an important application in archaeological studies, for example, analyses of ancient DNA may provide insights into the demographic structure, culture and social hierarchy of ancient individuals [5,13,14]. For nonhumans, studying patterns of kinship relationships among individuals in a local population is useful for characterizing mammalian social organization and population biology [6].

Many methods have been proposed for kinship inference, and they work mainly by estimating the proportion of the genome shared identical by descent (IBD) between individuals [1,15,16]. Of these methods, allele frequency-based approaches [17,18] are computationally efficient and are widely used to detect cryptic relatedness in large genomic data sets [19–21]. However, these methods can only accurately infer close relationships (generally up to third degree) [18,22] and have reduced accuracy in identifying distant relatives (e.g., sixth- and seventh-degree relatives) [23]. Furthermore, allele frequency-based approaches may be biased in admixed samples because of admixture linkage disequilibrium [24]. In contrast, the IBD segment-based methods generally have higher accuracy and can identify more distant relationships than those that rely on allele frequencies of independent markers [23]. The key idea behind IBD segment detection is haplotype frequency. If the frequency of a shared haplotype (in phased genotype data) or a half-shared segment (in unphased genotype data) is so small that it is unlikely to be observed twice in independently sampled individuals, one can infer the presence of an IBD segment [25].

In general, haplotype-based methods can detect short IBD segments with greater accuracy than genotype-based methods can [26]. However, haplotype-based methods can break up a long IBD segment into a series of shorter IBD segments if there are phasing errors [27], which can be further exacerbated by genotyping errors. For example, Turner et al. [28] showed that hap-ibd, a well-known haplotype-based method, failed to identify most IBD segments and assigned almost all samples as unrelated with error rates ≥1%. Furthermore, phasing often requires a genetic map, which may not be available for the dataset of interest. In contrast, genotype-based methods do not require phasing, and a higher level of genotyping errors can be acceptable [28].

Several excellent tools have been developed for the detection of IBD segments using unphased genotype data, such as IBIS [29], IBDseq [30], and TRUFFLE [31], which are fast and accurate for application to DNA samples of relatively high quality and quantity. However, these methods may be powerless for challenging materials, such as ancient human remains, samples from crime scenes, and noninvasively collected tissues from wild animals (e.g., feces, hair, and urine) [28,32,33]. DNA isolated from these samples is, in most cases, fragmented, low in quantity and/or mixed with nonhost DNA. For example, only ∼1% of DNA extracted from fecal-derived samples is endogenous to the donor animal [34] and ancient DNA libraries often contain <1% endogenous DNA, with the majority of sequencing capacity taken up by environmental DNA [35]. It is error-prone to generate genetic data from these poor-quality DNA samples. Ozga et al. [36] showed that the rates of allelic dropout ranged from 0.66% to 18.42% when generating genome-scale data by shotgun sequencing, and whole genome and exome capture sequencing using feces, urine, dentin, and dental calculus from wild eastern chimpanzees. If the input DNA is lower than 20pg, genotype errors may even exceed 20% when using SNP microarrays [37]. When the genotype error is low (<1%), existing IBD segment methods outperform allele frequency-based approaches (e.g., KING [18]) for close and distant relationship inference but they are less robust with an increasing genotype error (1–5%) [28].

Here, we proposed a method, clusIBD, which detects IBD segments by identifying clusters of low opposite-homozygote or mismatch regions between samples using unphased genetic data. We evaluated and compared the performance of clusIBD with IBIS, TRUFFLE and IBDseq using simulated families, artificial challenging materials and a five-generation family from chambered tombs in Early Neolithic Britain [13]. Our results demonstrated that clusIBD could robustly detect IBD segments from materials in various quality and quantity and may be a promising tool for forensic genetics and ancient DNA analyses.

## Materials and Methods

### clusIBD algorithm

The clusIBD algorithm is illustrated in Figure 1. It works on unphased bi-allelic single nucleotide polymorphisms (SNPs) with a minor allele frequency (MAF) above 0.05. All the markers are arranged by physical position, and are divided into non-overlapping windows with a fixed number of SNPs. We first consider regions where two individuals share only one haplotype, i.e., IBD1. We calculate the rate of opposite homozygous genotypes (R_ohg_), inside each window, and obtain a large number of R_ohg_ values, generally more than a thousand, for each pair of individuals. For unrelated individuals, the R_ohg_ values depend on the MAFs of the SNP markers used and can be estimated as 2*p*^2^*q*^2^ for each marker [18], where *p* and *q* are the frequencies of the reference and alternative alleles. In the absence of genotype errors and mutations, windows from IBD segments will all have R_ohg_ values of zero because they share at least one allele at each site. When genotype errors are introduced, the R_ohg_ values of IBD segments are no longer all zero but are expected to be smaller than those of non-IBD segments.

**Figure 1.**
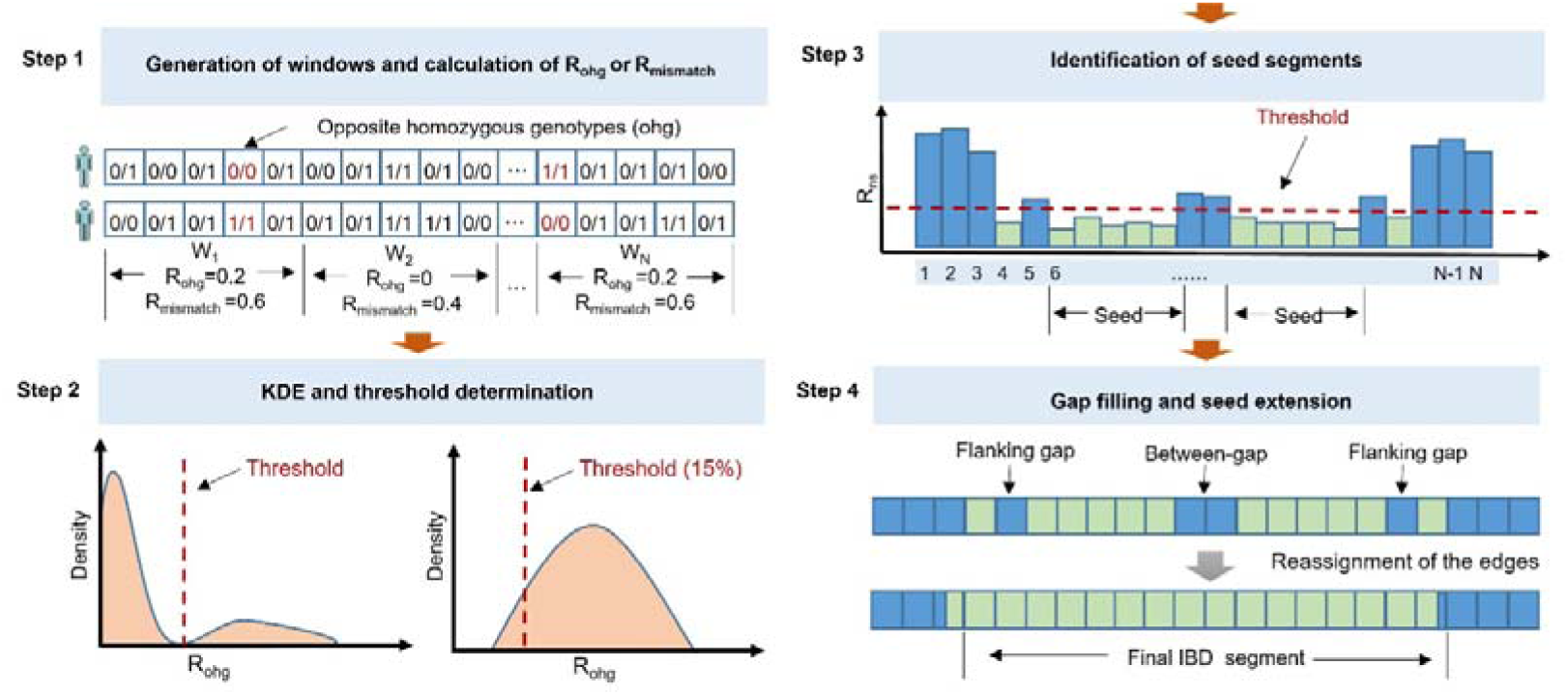
Illustration of clusIBD algorithm. Step 1: Obtain non-overlapping windows and calculate the rates of opposite homozygous genotypes (R_ohg_) for IBD1 or mismatch rates (R_mismatch_) for IBD2; Step 2: Perform Kernel density estimation and determine a threshold that may be used to distinguish windows from either IBD or non-IBD segments. Step 3: Identify consecutive windows all below the threshold and regard it as a seed segment. Step 4: Fill the gap between two seed segments and extend on its both sides if the flanking gaps are smaller than a predefined value. Finally, re-assign the start and end positions and report the IBD segments. IBD, identity by descent; IBD1, IBD in one copy of the genome; IBD2, IBD in both copies of the genome.

By leveraging this, a threshold (t) of R_ohg_ is determined and used to distinguish windows from either IBD or non-IBD segments. We first estimate the distribution of these R_ohg_ values using Kernal density estimation (KDE). If the two individuals have both IBD1 and non-IBD segments, we will see two peaks, the smaller one (near the y-axis) corresponding to windows of IBD segments (IBD peak) and the larger one corresponding to non-IBD segments (non-IBD peak; Figure 1). We will also see a valley between the two peaks, which is an ideal threshold (t_valley_) for distinguishing windows from either IBD or non-IBD segments. t_valley_ can then be determined using the R_ohg_ value corresponding to the inflection point, i.e., the point before which the probability density decreases and after which it increases. However, if the two individuals have unbalanced amounts of IBD and non-IBD segments, the IBD peak may be so small that it gets buried in the non-IBD peak and even becomes invisible, which is often observed at 4^th^ or more distant relationships (Supplementary Figure 1). To address this issue, we set a threshold corresponding to the 15% quantile (t_15th_) of the obtained R_ohg_ values for each pair of individuals with only one peak. Another problem arises when dealing with parent-child relationships, monozygotic twins, and genetic data from the same individual. Since they are homogeneous across the genome, we will also see only one peak and theoretically only 15% of the genome can be identified. To solve this problem, we randomly select several samples from the input data before the general analysis and calculate the heterozygote (*He*) rate for each window per sample. Since the expected *He* value is 2*pq*, a universal threshold (t_universal_) can be estimated as 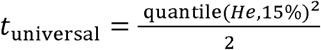. The final threshold is the maximum of t_valley_, t_15th_ and t_universal_. Because the threshold can be adapted to different levels of genotype error, clusIBD is always able to detect IBD segments at its best.

It is possible that some windows of non-IBD segments may have R_ohg_ values lower than the defined threshold. However, the probability of observing a region with a number of consecutive windows all below the threshold is very small. If such a region is identified, it is likely to be a part of an IBD segment and we call it a seed segment [26]. Due to stochastic effects, the R_ohg_ values of some windows of IBD segments may also be larger than the defined threshold, thus resulting in a series of interrupted segments. These gaps are addressed by using a gap-filling and seed-extension strategy. Specifically, if we identify two seed segments that are close to each other, we merge the two segments and the gaps between them into one long segment (i.e., gap-filling), considering that the probability of observing two recombination events within a small region is small. Similarly, if a long seed segment and a small gap on its either side are identified, we extend the boundary of the seed segment by skipping the gaps, i.e., seed extension. A similar process has also been applied in alternative tools [26,32]. This process is very useful for error control and makes clusIBD quite robust, especially when identifying IBD segments using genetic data with high genotype error (***Result****s*). Finally, the edges of a segment are reassigned, i.e., the last site with opposite homozygous genotypes in the first window as the final start position, and the first site with opposite homozygous genotypes in the last window as the final end position.

The same procedure can be used, with only some minor modifications, to infer the IBD2 segment. For IBD2 estimation, the rate of mismatches (R_mismatch_) or different genotypes, is calculate instead of R_ohg_. Since the main interference for IBD2 is not IBD0 but IBD1, we use a different procedure to determine the universal threshold 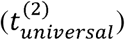. As IBD1 segments share one allele by descent at each site, the probability of having different genotypes is expected to be 2*pq*, i.e., *He*. Therefore, 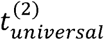 can be estimated as 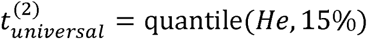.

### Implementation

We have implemented clusIBD in Python, including support for multithreading, and released it as open source (https://github.com/Ryan620/clusIBD/). The PLINK binary ped format (i.e., bed, bim, and fam files) is accepted as inputs. clusIBD outputs IBD segments in a human-readable text format. Two search modes are implemented, i.e. “within-search” and “across-search”. Within-search refers to inferring IBD segments for all the pairs within a database (one input file), whereas across-searching works by inferring IBD segments between two individuals from different databases (two input files). The latter is often performed when one or more individuals of interest are searched against a large database to identify potential relatives [38].

### Simulated data

Haplotype data of five representative populations from five major super populations, respectively, were extracted from the 1000 Genomes Project dataset (https://www.internationalgenome.org/). These populations are Utah Residents with North and West European ancestry (CEU) representing Europeans, Han Chinese in Beijing (CHB) representing East Asians, Gujarati Indian from Houston (GIH) representing South Asians, Mexican Ancestry from Los Angeles (MXL) representing admixed Americans, and Yoruba in Ibadan (YRI) representing Africans. For each population, we only included bi-allelic autosomal SNPs with a MAF greater than 0.05 and with a physical distance over 2 kilobases (kb). We then randomly selected a subset of these SNPs to create a reference panel of approximately 400,000 SNPs. Based on these panels, artificial IBD segments and family data were simulated. Specifically, we randomly selected a pair of individuals and a region of different sizes, and then a segment from one individual was injected into the same positions of another individual, thus generating an artificial IBD segment (IBD1). The phase state was discarded and four levels of genotype error were introduced, i.e., 0, 1%, 5%, and 10%. Pedigrees were simulated using Ped-sim [39] with sex-average genetic map [40]. From the simulated pedigree, we can obtain various relationships, including parent-child, full siblings, and 2^nd^ - 7^th^ degree relatives as well as unrelated pairs.

### Artificial poor-quality DNA

To test the performance of clusIBD on challenging materials, we generated a series of low-quantity and low-quality samples. Specifically, whole blood samples were collected with informed consents from six donors from a multi-generation family (Supplementary Figure 2). DNA was then extracted using the QIAamp DNA Blood Mini Kit (Qiagen, Hilden, Germany) and quantified on a Qubit® 3.0 fluorometer (Thermo Fisher Scientific, South San Francisco, CA) according to the manufacturer’s instructions. We further diluted these samples by adding nuclease-free water to mimic low-quantity DNA, and the final amounts of DNA were 100 nanograms (ng), 10 ng, 1 ng, 0.5 ng, and 0.1 ng for each sample. In addition, we fetched 100 ng of DNA and degraded them using the Covaris M220 focused ultrasonicator (Covaris, Inc.). By controlling the duty factor and treatment time, four levels of degraded DNA were obtained, with fragment sizes of approximately 2500 base pairs (bp), 800 bp, 400 bp, and 150 bp, respectively. These samples were finally genotyped using the Infinium Asian Screening Array (ASA; Illumina Inc., San Diego CA). SNPs with a MAF of less than 0.05, or on a non-autosomal chromosome were excluded, thus resulting in approximately 300 thousand SNPs. Genetic analysis of the family was approved by the Ethics Committee of Zhongshan School of Medicine, Sun Yat-sen University (Guangzhou, China), approval number [2023]016.

### Ancient DNA

Thirty-five individuals who lived about 5,700 years ago and were entombed at Hazleton North long cairn were deeply characterized by combining archaeological and genetic data in Fowler et al’s study [13]. Pedigree reconstruction indicated that these samples consisted of twenty-seven individuals from a five-generation lineage and eight unrelated individuals. We downloaded the aligned sequencing data (bam files) from the European Nucleotide Archive with accession number PRJEB46958. SNP calling was performed at 1,233,013 sites of the 1240K panel using Samtools [41]. We also downloaded genetic data of present-day individuals from England from the 1000 Genome Project dataset. We then removed SNPs with more than 50% missing calls and MAF less than 0.05, and individuals with more than 50% missing sites. With these filtrations, the final set consisted of 30 individuals genotyped at 361,399 SNPs.

### Relatedness Classification

For each pair of individuals, we calculated the kinship coefficient (θ) [42], which denotes the probability that a randomly selected allele in one individual is IBD with a randomly selected allele from the same genomic position of another individual. θ can be calculated as 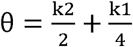, where k2 and k1 denote the proportions of the genomes that two individuals share IBD2, and IBD1, respectively. Thus, with reported IBD segments, θ can also be simply estimated as 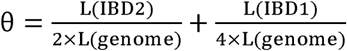, where L(IBD1), L(IBD2), and L(genome) are the total lengths of IBD1 segments, IBD2 segments, and the whole genome, respectively. The expected values of θ for different relationships are listed in Table 1. Following a decision-making process similar to KING [18], we inferred the degree of relatedness for a pair of individuals using the criteria in Table 1, and pairs with 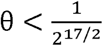 were assigned as unrelated.

**Table 1.**
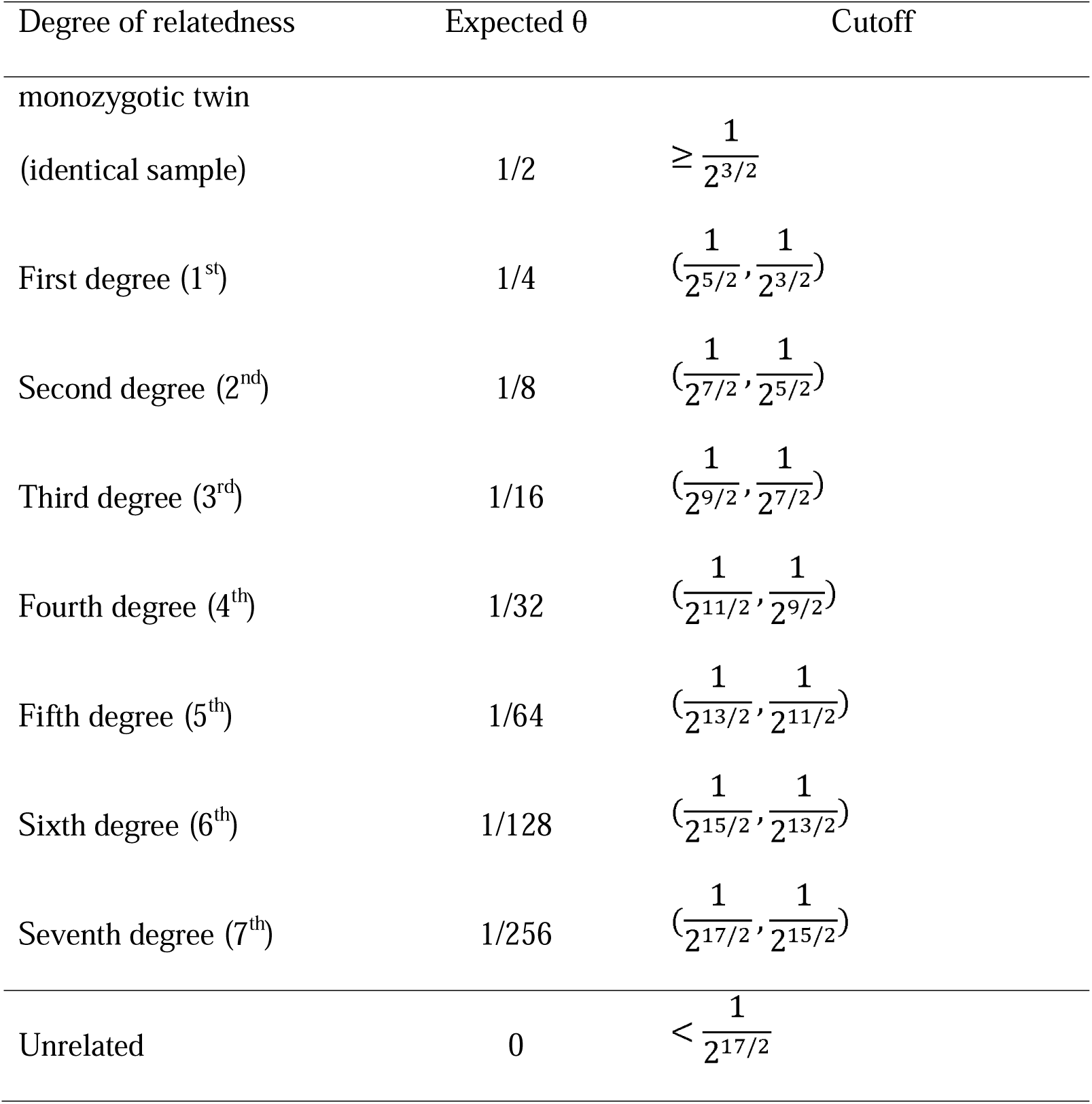
The expected values of kinship coefficients and inference criteria for different relationships.

| Degree of relatedness | Expected $\theta$ | Cutoff |
| --- | --- | --- |
| monozygotic twin |  |  |
| (identical sample) | 1/2 | $\geq \frac{1}{2^{3/2}}$ |
| First degree (1<sup>st</sup>) | 1/4 | $(\frac{1}{2^{5/2}}, \frac{1}{2^{3/2}})$ |
| Second degree (2<sup>nd</sup>) | 1/8 | $(\frac{1}{2^{7/2}}, \frac{1}{2^{5/2}})$ |
| Third degree (3<sup>rd</sup>) | 1/16 | $(\frac{1}{2^{9/2}}, \frac{1}{2^{7/2}})$ |
| Fourth degree (4<sup>th</sup>) | 1/32 | $(\frac{1}{2^{11/2}}, \frac{1}{2^{9/2}})$ |
| Fifth degree (5<sup>th</sup>) | 1/64 | $(\frac{1}{2^{13/2}}, \frac{1}{2^{11/2}})$ |
| Sixth degree (6<sup>th</sup>) | 1/128 | $(\frac{1}{2^{15/2}}, \frac{1}{2^{13/2}})$ |
| Seventh degree (7<sup>th</sup>) | 1/256 | $(\frac{1}{2^{17/2}}, \frac{1}{2^{15/2}})$ |
| Unrelated | 0 | $< \frac{1}{2^{17/2}}$ |

## Results

### IBD segment detection

First, we evaluated the performance of clusIBD in detecting IBD segments of different lengths, ranging from 10 megabases (Mb) to 30 Mb, and with different levels of genotype error, using simulated IBD segments. It is worth mentioning that genetic distance is often a better unit to use for length-based IBD detection, but a genetic map may not be available for the dataset of interest (e.g., ancient DNA samples). Based on our established reference panels, five hundred artificial IBD segments were simulated for each length and error group as described in *Material and Methods*. Since we know the true breakpoints of each ground-truth IBD segment, we can compare them to the reported IBD segments to evaluate the performance. However, the comparison is not straightforward because the reported segments may be partially true, or a ground-truth IBD segment may be partially reported. Therefore, we used the four metrics defined by Tang et al. [43], including recall and power to evaluate the ground-truth IBD segments, and accuracy and length accuracy (len.accuracy) to evaluate the reported IBD segments. Additionally, we also compared the performance with the other three methods, i.e., IBIS, TRUFFLE, and IBDseq. The default options for all four methods were used throughout the analyses unless otherwise specified.

The results showed that clusIBD was accurate down to 7 Mb, whereas for shorter IBD segments, a large proportion of identified segments were false positives (Supplementary Figure 3), which has also been previously reported for alternative methods [29,31,44]. To eliminate potential spurious signals, segments smaller than 7 Mb were excluded for all the four methods. Figure 2 shows the performance of clusIBD and three alternative methods (IBIS, TRUFFLE, and IBDseq) for detecting IBD segments. The four methods performed similarly when the genotype error was low. However, clusIBD was much more robust when the genotype error was above 1%. For example, at an error rate of 5%, approximately 85% of the 10 Mb segments can be recalled with clusIBD, while the recall rates for the other three methods were all less than 10%. It appears that clusIBD had a higher power to detect long IBD segments than the short ones. When the genotype error exceeded 10%, IBIS, TRUFFLE and IBDseq recalled almost no segments. Notably, clusIBD was still able to detect many IBD segments at such high error rates. More importantly, when we used a larger number of SNPs per window, more segments can be detected with a higher accuracy but at the cost of decreasing call rates for short segments (Supplementary Figure 4).

**Figure 2.**
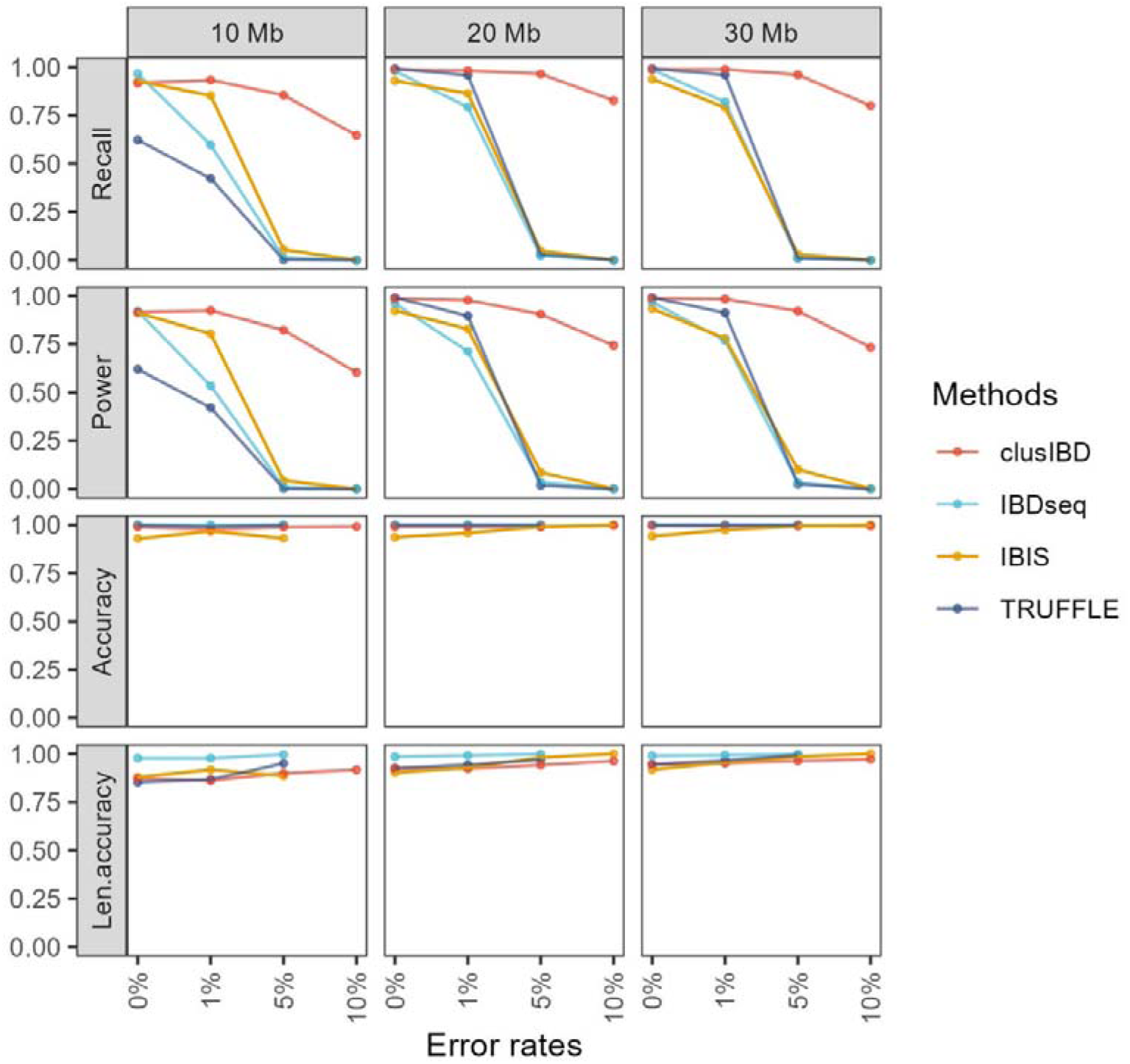
Performance of clusIBD, IBDseq, IBIS, and TRUFFLE for detecting IBD segments of different lengths and with different levels of genotype error. Recall is the proportion of ground-truth IBD segments that could be covered by any one reported IBD segment by >50%. Power measures the proportion of lengths of ground-truth IBD segments that are covered by a best-reported IBD segment. Accuracy is the proportion of reported IBD segments that are covered by any one ground-truth IBD segment by >50%. Len.accuracy is the proportion of the maximum lengths overlapped between the ground-truth and detected IBD segment divided by the reported lengths. For details, please refer to Tang et al, Gigascience, 2022 [43]. IBD, identity by descent.

In terms of the accuracy of IBD segments detected, clusIBD, IBDseq and TRUFFLE showed similar accuracies, slightly higher than IBIS (Figure 2 and Supplementary Figure 5). IBDseq performed the best among the four methods, with almost 100% accuracy in detecting IBD segments of different lengths when the genotype error was no more than 5% (Supplementary Figure 5). However, when the error rate exceeded 5%, IBDseq could only detect one or two segments for each length group, most of which were marginally larger than the predefined threshold, and some of them were incorrect. In contrast, clusIBD could detect many more IBD segments with a consistent accuracy for genetic data with different levels of genotype error. When the lengths were taken into consideration for accuracy estimation (i.e., len.accuracy), IBDseq performed the best, and clusIBD, IBIS, and TRUFFLE showed similar len.accuracy (Figure 2 and Supplementary Figure 5). Due to random matches at SNP sites of non-IBD segments, clusIBD (also IBIS and TRUFFLE) tended to underestimate the start positions and overestimate the end positions (Supplementary Figure 6). Nevertheless, the median difference between estimated and actual breakpoints was less than 0.5 Mb and the difference decreased with increasing genotype error, resulting in a slightly higher len.accuracy. See Supplementary Figure 6 and Supplementary Table 1 for details.

Since our method is a data-driven approach, we further investigated whether the detection of IBD segments varied for different relationships and for samples from different populations. The ground truth IBD segments of simulated 1^st^ - 7^th^ degree relationships were categorized into three segment size bins: [7 Mb, 15 Mb), [15 Mb, 25 Mb) and [25 Mb, 35 Mb) and segments larger than 35 Mb were not included. The results showed that there was a slight but consistent decrease in recall and power to detect IBD from distant relationships, which was more evident for shorter segments and exacerbated by higher genotype error (Supplementary Figure 7). Surprisingly, a slight decrease in recall and accuracy was also observed for first-degree relationships. This can be explained by the fact that some adjacent ground-true but short segments were merged into one long segment by clusIBD. As a result, the reported segments were not considered correct because they were not covered by ≥50% by any ground-truth IBD segment. In addition, clusIBD performed very similarly for samples from different populations, except for a slight decrease for Mexican ancestry from Los Angeles (MXL; Supplementary Figure 8).

### Kinship inference of simulated data

We then evaluated the performance of clusIBD in classifying relationships ranging from first- to seventh-degree and unrelated individuals using simulated family data based on the 1000 Genomes Project dataset. IBDseq only reports IBD segments, regardless of whether they are IBD1 or IBD2. Since a pair of full siblings is expected to share 25%, 50% and 25% of IBD2, IBD1 and non-IBD segments, respectively, thus two thirds of these reported segments are expected to be IBD1 and the remaining third to be IBD2. Therefore, we estimated the total length of IBD1 by multiplying the reported IBD by 2/3 and for IBD2 by 1/3 for a full sibling pair. Kinship coefficient estimation and relationship classification were performed as described in ***Material and Methods*.**

In the absence of genotype error, the agreement of kinship coefficients between clusIBD and IBIS, TRUFFLE, and IBDseq was very high, with R^2^ equal to 0.999, 0.999, and 0.997, respectively (Supplementary Figure 9). IBIS tended to slightly underestimate the kinship coefficients for full siblings compared to the other three algorithms at error rates of 1%. After reviewing the results of IBIS, we found that many IBD2 segments were misclassified as IBD1 segments, explaining the decrease in kinship coefficients for full siblings with this method. Furthermore, R^2^ values dropped dramatically at an error rate of 5% and kinship coefficients were all largely underestimated with IBIS, TRUFFLE, and IBDseq. In contrast, clusIBD could robustly estimate kinship coefficients at an error rate as high as 10%. In terms of relationship classification, all algorithms performed well and had comparable recall rates for inferring close and distant relationships from genetic data with no genotype error (Figure 3). clusIBD outperformed IBIS, TRUFFLE, and IBDseq when the genotype error was 1% or higher. In particular, when the three alternative methods showed near-zero recall rates at 5% error for all the relationships, clusIBD still successfully recalled them with high accuracy. Notably, IBIS had low power to identify full siblings even at a low level of genotype error (1%), which may be explained by its low power to identify IBD2 segments in the presence of genotype error. These results demonstrate that clusIBD can robustly infer a relationship from genetic data with varying levels of genotype error and outperforms the three alternative methods.

**Figure 3.**
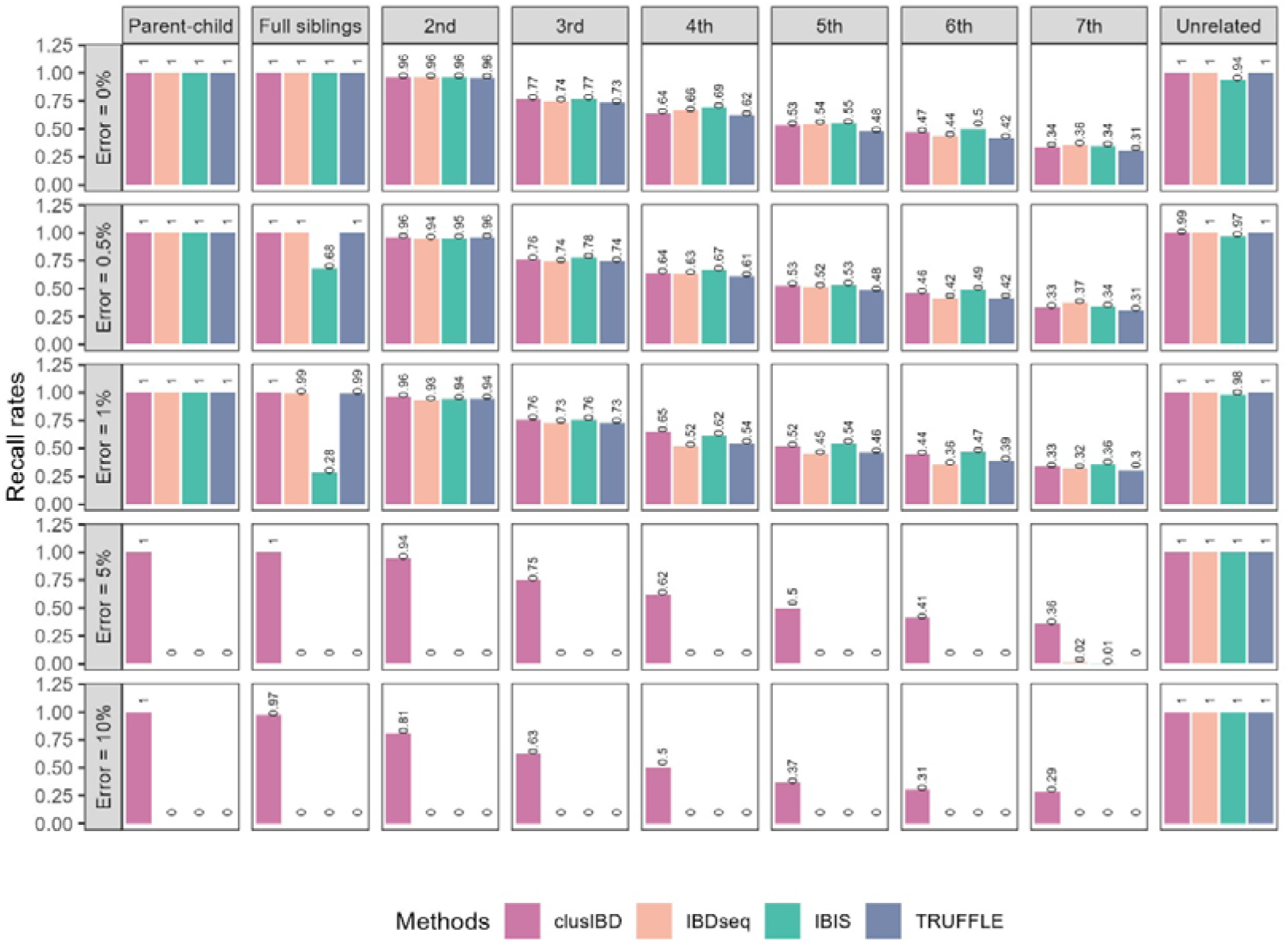
Comparison of clusIBD, IBDseq, IBIS, and TRUFFLE in kinship inference at different levels of genotype error. Recall rate is the proportion that a relationship is correctly assigned as the true relationship.

### Kinship inference of poor-quality DNA

DNA from six individuals from a multi-generation family (Supplementary Figure 2) was collected with informed consent (Human Subject IRB Protocol identifier: [2023]016, approved by the Ethics Committee of Zhongshan School of Medicine, Sun Yat-sen University), and was then diluted and degraded to mimic low-quantity and low-quality samples. These samples were finally genotyped using the Infinium Asian Screening Array (ASA; Illumina Inc., San Diego CA). After quality control and SNP filtering (see *Material and method*), approximately 300 thousand SNPs were retained. Prior to IBD detection, we evaluated the level of genotype error for these artificial challenging DNA samples. For each sample, the 100 ng non-degraded DNA group was treated as a reference, and three types of genotype error (drop-in, dropout, and opposite-homozygote errors) were estimated for the remaining samples. Drop-in error refers to a homozygote being reported as a heterozygote, while dropout refers to a heterozygote being reported as a homozygote; opposite-homozygote error refers to a scenario where different homozygotes are reported. As shown in Supplementary Figure 10 and Supplementary Table 2, although 200 ng of high-quality DNA is recommended by the manufacturer, we found that DNA as low as 0.5 ng or with a fragment size of approximately 800 bp still had low genotype errors (error rates < 0.5%). However, when the DNA input was very low (0.1 ng) or highly degraded (fragment size of approximately 150 bp and 400 bp), genotype errors increased significantly, with average error rates of 4.30% ± 2.36%, 12.49% ± 1.24%, and 0.84% ± 0.26% for the 0.1 ng, 150 bp, and 400 bp groups, respectively. It appears that the pattern of genotype errors differed between different groups. The predominant error was allele dropout for the low-quantity group (i.e., 0.1 ng group), whereas it was allele drop-in for the low-quality groups (i.e., 150 bp and 400 bp groups). The rates of opposite-homozygote error were extremely low for all groups (Supplementary Figure 10 and Supplementary Table 2).

For kinship inference, we mainly focused on the low-quantity and low-quality groups, including the 0.1 ng, 150 bp, and 400 bp groups. With the 18 samples (six samples for each group), we had 36, 18, 9, 18, 27, 18 and 9 pairs of 1^st^ to 7^th^ degree relationships, respectively, as well as 18 pairs of identical samples (Figure 4 and Supplementary Figure 2). We then used the four tools to detect IBD segments in the dataset, and the relationship was determined as described above. Overall, clusIBD had the highest overall accuracy of 31.37% compared to TRUFFLE (18.30%), IBIS (8.50%), and IBDseq (0%). IBDseq detects IBD segments based on IBD LOD scores, which is calculated using the variant’s MAF. It also requires LD-based thinning of the variants so that no pair of variants is strongly correlated [30]. However, all the samples in this dataset were related to each other, and the MAF estimation and LD-based thinning cannot be handled correctly, which may explain the poor performance of IBDseq. For each type of relationship, the recall rates were 100% (18/18), 41.67% (15/36), 44.44% (8/18), 44.44% (4/9), 16.67% (3/18), 0%, 0%, and 0% for identical individuals and 1^st^ to 7^th^ degree relationships, respectively, all higher than the rates with the other three methods. These values increased to 100%, 63.89%, 61.11%, 66.67%, 38.89%, 18.52%, 16.67%, and 0% if one degree difference was allowed. Although the nine pairs of 7^th^ degree relationships were all incorrectly assigned as unrelated, a long IBD segment of 16 - 22 Mb was identified on chromosome 1 for six out of the nine pairs (Supplementary Table 3), strongly suggesting their relatedness.

**Figure 4.**
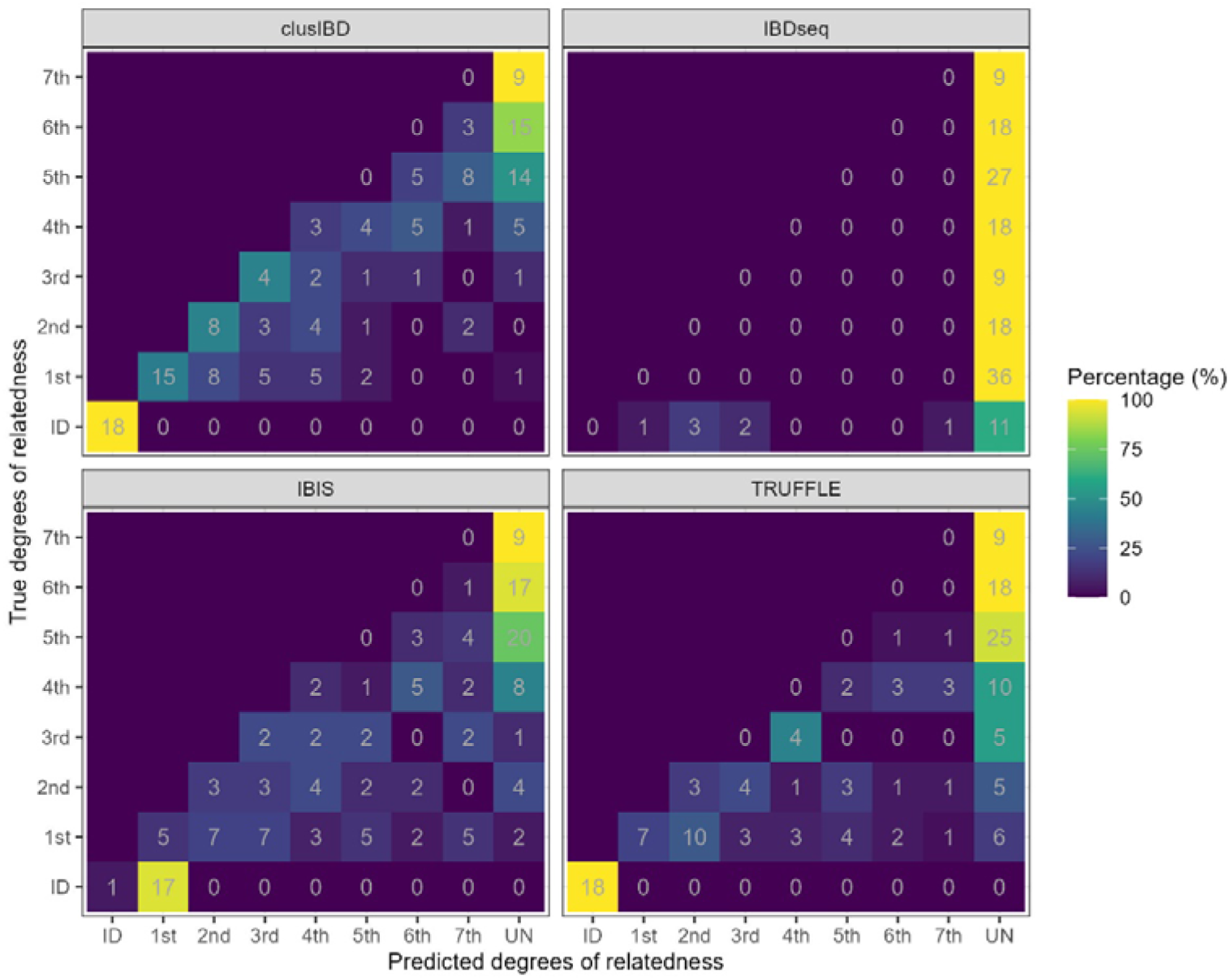
Comparison of clusIBD, IBDseq, IBIS, and TRUFFLE in kinship inference for poor-quality DNA samples. Eighteen samples were included from the 0.1 ng, 150 bp, and 400 bp groups. In total, 36, 18, 9, 18, 27, 18 and 9 pairs were obtained for 1^st^ to 7^th^ degree relationships, respectively, as well as 18 pairs of identical samples. ID, identical samples; UN, unrelated individuals; ng, nanograms; bp, base pairs.

### Kinship inference of ancient DNA

Finally, we analyzed the thirty ancient samples from the Hazleton North long cairn. The ancient DNA was sequenced by Fowler et al [13] using whole genome capture technology. One sample (SC5m) was removed because it could not be explicitly positioned in the pedigree by the authors of the original study. As shown in Figure 5, clusIBD reported larger total IBD lengths than IBIS, TRUFFLE, and IBDseq for the majority of related individuals, indicating the high robustness of clusIBD to genotype error. Of the three alternative tools, TRUFFLE, which incorporates an error model corrects for segment break-ups that occur as a consequence of genotype errors [31], performed the best while IBDseq showed the worst performance. We further inferred the relationships as described in ***Material and Methods*** and compared the results to the pedigree constructed by Fowler et al [13]. With the default settings, the estimated number of SNPs per window was 150. Only one pair of related individuals was correctly assigned, resulting in a concordance rate of 0.50% (Supplementary Figure 11A and 11B), which increased to 3.02% with 300 SNPs per window (Supplementary Figure 11C and 11D), and 14.07% with 500 SNPs per window (Figure 5B), approximately six and thirty times higher than with the default settings, respectively. More interestingly, although the relatedness was underestimated for most of these individuals, four pairs that were once recognized as unrelated in the study by Fowler et al. were inferred as 7^th^ degree relationships with clusIBD. The four pairs involved four family members and two unrelated individuals. NE4m, one of the five unrelated males inferred by Fowler et al., shared a long segment (> 30Mb) with three family members, i.e., SC2m, SP1m, and SE1m (Supplementary Table 4). SC2m and SP1m were full siblings and both shared an approximately 36 Mb segment with NE4m at almost the same positions (chr3: 80 - 117 Mb). NE4m also shared a 32 Mb IBD segment at chr14: 51 - 83 Mb with SE1m, who was the offspring of a distant relative of the core family member NC1m. We speculated that NE4m was a descendant of a distant relative of NC1m, considering that the introduction of non-paternal male into the tomb was rare. Similarly, NC10m also shared a long segment with two family members (i.e., SP3m and NC7f, who were also full siblings; Supplementary Table 4) and might be distantly related to the family. In summary, clusIBD not only successfully recovered some of the related individuals from the thirty ancient samples but also identified two additional males as related to the family. Since only three males in the grave were unrelated to the family, we may conclude that it is rare for kinship in this period to include social ties independent of biological relatedness.

**Figure 5.**
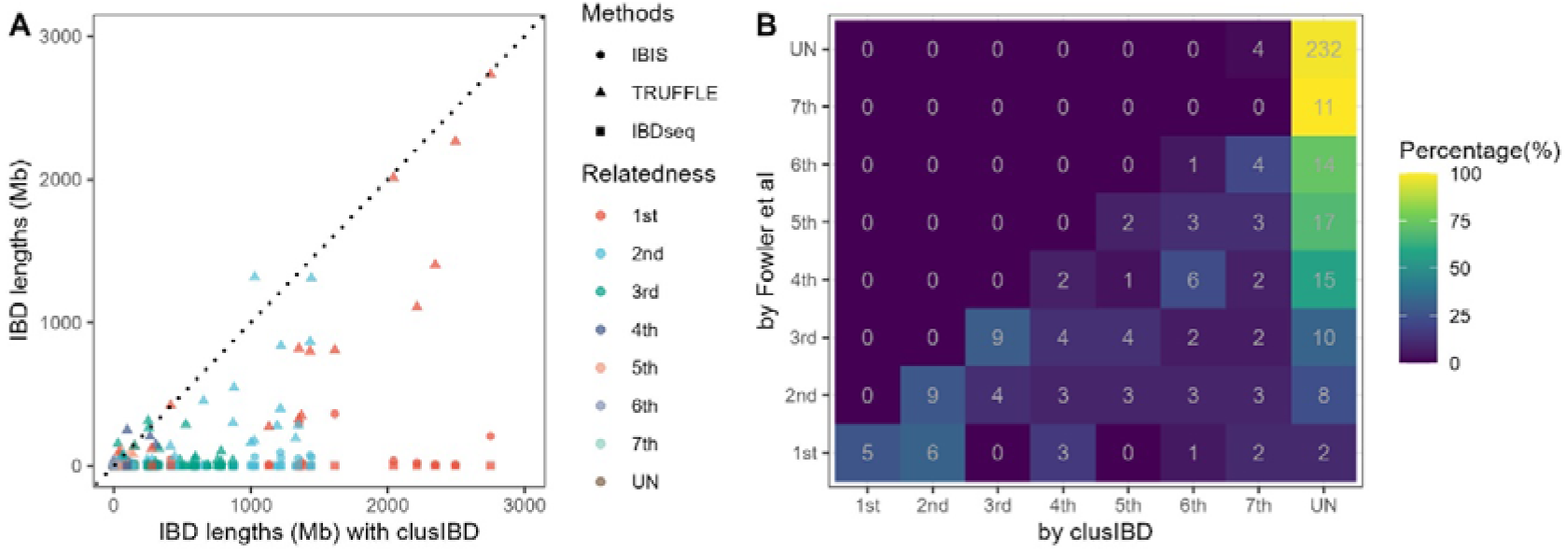
IBD lengths and kinship inference for ancient DNA samples. (A) Scatter plots of IBD lengths estimated by clusIBD and IBIS, TRUFFLE, and IBDseq. Dotted lines represent a line with a slope of 1, i.e., y=x, and points below the line indicate a larger IBD lengths by clusIBD than by IBIS (circle), TRUFFLE (triangle), and IBDseq (square). (B) Match matrices of inferred relationships by Fowler et al. and by clusIBD. 500 SNPs per window was used. SNPs, single nucleotide polymorphisms; IBD, identity by descent; UN, unrelated individuals.

### Computation time

Five datasets with sample sizes of 100, 200, 300, 400 and 500 individuals were simulated based on our established reference panels and the computing time of clusIBD, IBIS, TRUFFLE and IBDseq were compared. Since IBDseq only analyzes one chromosome at a time, the computing time was estimated by simply summing the time over 22 chromosomes. All the four tools allow for parallel computation with a specified number of cores, and we reported the time using an Intel(R) Xeon(R) CPU E5-2650 v4 @ 2.20GHz processor with five cores. For the 100-sample batch, clusIBD ran in 47 seconds (s), and the computing time increased quadratically as the sample size increased (Supplementary Figure 12). For batches of size 500 samples, the average time cost was about 0.03 s per core per pair of samples. IBIS was the most computationally efficient algorithm, taking only about 2 s for 100 samples. In contrast, IBDseq was the least efficient, requiring more than 200 s (100 samples). The main time cost was the LD-based thinning for IBDseq. Overall, clusIBD was faster than IBDseq but slower than IBIS and TRUFFLE.

## Discussion

We have proposed a method, clusIBD, for detecting IBD segments using unphased genetic data with different levels of genotype error, and we have compared clusIBD with several existing IBD detection methods using simulated family data and real SNP array and whole genome sequencing data. Our results demonstrated that clusIBD can robustly detect IBD segments from materials of varying quality and quantity, and may be a promising tool for forensic genetics and ancient DNA analysis.

One of the key advantages of our method is its robustness to genotype error. Existing IBD detection methods are fast and accurate when applied to samples of high quality and quantity, but they only allow relatively low levels of genotype errors (0.1% - 1% by default) [29–31]. Many materials, such as ancient DNA, crime scene samples, and noninvasively collected tissues [36,37], are error-prone and typically have genotype errors above this limit, rendering them incapable of detecting IBD segments from these samples. In contrast, clusIBD works by determining a threshold to distinguish IBD segments from non-IBD segments based on the difference in R_oph_ distributions.

As the thresholds can be adapted to the level of genotype error for each sample pair of interest, clusIBD was always able to detect IBD segments at its best. In addition, for extremely challenging materials, users can further improve the robustness by increasing the number of SNPs per window, regardless of the decrease power in identification of short IBD segments (Supplementary Figure 4). Although IBIS also allows users to increase the acceptable error rate in a segment before considering it false, with the goal of identifying more IBD segments in the case of genotype error, however, the method is not an error-aware algorithm and increasing the threshold results in a reduction in accuracy (Supplementary Figure 13). Turner et al. [28] showed that there was no combination of reasonably permissive parameters that could rescue the performance of the IBD segment methods they studied (e.g., IBIS) when the genotype error was in the 1 - 5% range. Therefore, it may be impractical to simply increase the acceptable error rates, which increases the risk of false positives. Another way to control genotyping error is the use of gap-filling and seed-extension strategy. A similar process has also been applied in alternative tools for phased data [26,32]. Freyman et al. [27] show that some IBD detection methods create short but very close together IBD segments, which are in fact small parts of a long single long true IBD segment and should be stitched together. In view of this, some methods also try to address genotyping errors post hoc, e.g., ibd-end [45] and IBDkin [16]. All of these processes are very useful for improving the robustness of related methods to genotyping errors. Some DTC companies (e.g., FTDNA and 23andMe) and third-party services (e.g., GEDmatch) use similar half-identical matching algorithms to detect IBD segments with unphased genotypes [44]. We uploaded the genotype files of the eighteen poor-quality DNA samples (0.1 ng, 150 bp, and 400 bp groups) to GEDmatch, a well-known third-party service and the main entry point for law enforcement up till now. We found that clusIBD recalled longer total IBD segments for most of the sample pairs than GEDmatch, thus outperforming the latter (Supplementary Figure 14).

clusIBD uses a simple window-based approach, and identifies IBD segments by searching successive windows that are all below a predefined threshold (R_ohg_ for IBD1 and R_mismatch_ for IBD2), making it more computationally efficient than likelihood and hidden Markov model (HMM)-based methods [30,32]. clusIBD can be easily used without complex data pre-processing, such as haplotype phasing, which is time-consuming and has a risk of phasing errors [23,43,46]. Nevertheless, IBD detection can be performed quite efficiently with phased data [26,27,47–50].

The pseudohaploid-based approach is also computationally efficient and robust to genotype error, and has been widely applied to the analysis of ancient DNA [51]. Pseudohaploid genotype calling allows for genetic information to be obtained at a nucleotide site covered by only a single read. Several tools have been developed [22,52,53] and kinship inference can be successfully performed with as little as 0.1x shotgun sequencing data [22]. However, these methods either require for a dataset of sufficient sample size to provide a reliable normalization value or require the user to provide a set of allele frequencies, which can be challenging for unknown forensic human remains, archaeological samples, and small conservation populations [54]. More importantly, if the allele frequencies used are not representative of the target population, there will be significant overrepresentation of the kin relationships [55]. In addition, similar to the frequency-based approaches, pseudohaploid-based approaches can only accurately infer close relationships and have high false-positive rates in identifying distant relatives.

For relationship classification, IBD segment-based methods have high accuracy and can identify very distant relationships [23]. We can also discriminate relationships of the same degree of relatedness based on different IBD signals [56,57]. Qiao et al. showed that specific pedigree relationship types could be uncovered via multi-way IBD sharing, and they were able to classify pairs as maternally or paternally related with sex-specific genetic map [58]. In addition to relatedness classification, IBD segments are also useful for a variety of analyses, including studies of population history [59], estimation of mutation rates [60] and recombination rates [61], disease gene mapping [50,62–64], and recent positive selection [45,65].

There may be some limitations of our method. clusIBD has reduced sensitivity in identifying short IBD segments, which may be acceptable for certain analyses because recently related individuals are expected to share long IBD segments. In addition, there is potential to overcome this by allowing for more SNP sites to be analyzed. Another limitation of our approach is the inability to infer IBD segments on the X chromosome, which is useful to study sex-specific migration patterns [66].

## Conclusions

In summary, clusIBD can robustly detect IBD segments from materials of varying quality and quantity using either SNP array or whole genome sequencing data. It can be used to accurately infer relationships from unphased genetic data with high genotype errors (∼10%) and is a promising tool for forensic genetics and ancient DNA analysis.

## CRediT author statement

**Ran Li**: Conceptualization, Methodology, software, Writing-Origin draft, Funding acquisition. **Yu Zang**: Investigation, Formal analysis, Data curation, Writing - Review & Editing. **Zhentang Liu**: Methodology, software. **Jingyi Yang**: Investigation, Validation. **Nana Wang**: Visualization, Validation. **Jiajun Liu**: Visualization. **Enlin Wu**: Investigation. **Riga Wu**: Supervision, Project administration. **Hongyu Sun**: Conceptualization, Writing - Review & Editing, Project administration, Funding acquisition. All authors contributed to the final text and approved it.

## Acknowledgements

The authors would like to thank all the volunteers. This research was funded by the National Natural Science Foundation of China under Grant number 82302125 and by the National Key R&D Program of China under Grant number 2024YFC3306702.

## Availability of data and materials

The software used in this manuscript can be downloaded from the appropriate repositories:

Ped-sim, https://github.com/williamslab/ped-sim;

TRUFFLE, https://adimitromanolakis.github.io/truffle-website/;

IBDseq, http://faculty.washington.edu/browning/ibdseq.html;

IBIS, https://github.com/williamslab/ibis/; Our software is also available in the GitHub repository (https://github.com/Ryan620/clusIBD/).

The raw data of the thirty-five ancient DNA samples can be accessed at the European Nucleotide Archive under the accession number PRJEB46958 (https://www.ebi.ac.uk/ena/browser/view/PRJEB46958). We deposited the haplotype data of 208 Chinese individuals from the 1000 Genomes Project dataset, which were used for family simulation and IBD segment generation (https://github.com/Ryan620/clusIBD/tree/main/source_code/0.referenceData). The genetic data of the six modern family members that support the finding of this study are not publicly available due to privacy concerns. However, it is available upon request from the corresponding author (Hongyu Sun). All the source codes for data analysis and figure generation are also available in the GitHub repository (https://github.com/Ryan620/clusIBD/). Access to the data requires noncommercial, research only usage approved by the ethical committee.

## Declaration of generative AI and AI-assisted technologies in the writing process

During the preparation of this work the author(s) used DeepL (https://www.deepl.com/) in order to improve readability and language. After using this tool/service, the author(s) reviewed and edited the content as needed and take(s) full responsibility for the content of the publication.

## Supplementary materials

### Supplementary Tables

**Supplementary Table 1. The median difference (Mb) between estimated and actual breakpoints by clusIBD**

Note: A negative value indicates an underestimated position and a positive value indicates an overestimated position. Mb, megabases.

**Supplementary Table 2. The types and rates of genotype error of the artificial poor-quality DNA samples**

Note: Drop-in error refers to a homozygote being reported as a heterozygote, while dropout refers to a heterozygote being reported as a homozygote; opposite-homozygote error refers to a scenario where different homozygotes are reported. ng, nanograms; bp, base pairs.

**Supplementary Table 3. IBD segments identified by clusIBD for the artificial poor-quality DNA samples**

Note: IBD, identity by descent; IBD1, IBD in one copy of the genome; IBD2, IBD in both copies of the genome; ng, nanograms; bp, base pairs.

**Supplementary Table 4. IBD segments identified by clusIBD for the ancient DNA samples**

Note: IBD, identity by descent; IBD1, IBD in one copy of the genome; IBD2, IBD in both copies of the genome.

### Supplementary Figures

**Supplementary Figure 1. Distributions of the rates of opposite homozygous genotypes for different relationships**

In this example, we show a scenario where one person has an error rate of 10%, while many others (including relatives of the person of interest) in the dataset have ∼0.5%. Bimodal peaks are observed for 1^st^ - 3^rd^ degree relationships, the smaller of which (i.e. near the y-axis) corresponds to windows of IBD segments and the larger of which corresponds to non-IBD segments. The three metrices, i.e., t_universal_, t_valley_ and t_15th_, are estimated by clusIBD and the final threshold is the maximum of them. For close relationships, t_valley_ is greater than t_universal_ and t_15th_, and so would be the threshold. However, for distant (4^th^ degree or more distant) relationships, null results are obtained for t_valley_, and t_universal_ is greater than t_15th_, so t_universal_ is chosen. Sometimes, t_15th_ is greater than t_universal_ for very distant relationships and unrelated individuals, and t_15th_ is used as the threshold. Since the threshold can be adapted to different levels of genotype error for the sample pair of interest, clusIBD is always able to detect IBD segments at its best. IBD, identity by descent.

**Supplementary Figure 2. The pedigree of the studied family**

Samples filled in blue were collected.

**Supplementary Figure 3. Performance of clusIBD on the detection of IBD lengths ranging from 2 Mb to 15 Mb**

A total of five hundred artificial IBD segments are analyzed for each length group. IBD, identity by descent; Mb, megabases.

**Supplementary Figure 4. Performance of clusIBD with different numbers of SNPs per window**

A total of five hundred artificial IBD segments are analyzed for each length and error group. IBD segments of 20 Mb in length are used for parts A, B and C. SNPs, single nucleotide polymorphisms; IBD, identity by descent; Mb, megabases.

**Supplementary Figure 5. Performance of clusIBD, IBIS, TRUFFLE, and IBDseq for detecting IBD segments (zooming in on the 0.75 – 1.00 y-axis region)**

Accuracy is the proportion of reported IBD segments that are covered by any one ground-truth IBD segment by >50%. Len.accuracy is the proportion of the maximum lengths overlapped between the ground-truth and detected IBD segment divided by the reported lengths. For details, please refer to Tang et al, Gigascience, 2022 [43]. IBD, identity by descent; Mb, megabases.

**Supplementary Figure 6. The distribution of the difference between estimated and actual breakpoints by clusIBD**

A negative value indicates an underestimated position and a positive value indicates an overestimated position. Mb, megabases.

**Supplementary Figure 7. Performance of clusIBD for detecting IBD segments from different relationships**

The ground truth IBD segments for 1^st^ to 7^th^ degree relationships are categorized into three segment size bins: [7 Mb, 15 Mb), [15 Mb, 25 Mb) and [25 Mb, 35 Mb) and segments larger than 35 Mb were not included. IBD, identity by descent; Mb, megabases.

**Supplementary Figure 8. Performance of clusIBD for detecting IBD segments between samples from different populations**

CEU, Utah Residents with North and West European ancestry; CHB, Han Chinese in Beijing; GIH, Gujarati Indian from Houston; MXL, Mexican Ancestry from Los Angeles; and YRI, Yoruba in Ibadan. IBD, identity by descent; Mb, megabases.

**Supplementary Figure 9. Scatter plots of kinship coefficients (**θ**) estimated by clusIBD and IBIS, TRUFFLE, and IBDseq**

The θ values are estimated using the formula: 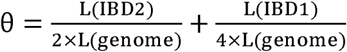, where L(IBD1), L(IBD2), and L(genome) are the total lengths of IBD1 segments, IBD2 segments, and the whole genome, respectively. Since IBDseq does not distinguish between IBD1 and IBD2 segments, we estimated the total length of IBD1 by multiplying the reported IBD by 2/3 and for IBD2 by 1/3 for a full sibling pair. The dotted black lines represent a line with a slope of 1, i.e., y=x. IBD, identity by descent; IBD1, IBD in one copy of the genome; IBD2, IBD in both copies of the genome.

**Supplementary Figure 10. The rates of genotype errors of the artificial poor-quality DNA**

Drop-in, a homozygote is reported as a heterozygote; dropout, a heterozygote is reported as a homozygote; opposite-homozygote error, opposite homozygotes are reported; all, the sums of the three types of genotype errors; ng, nanograms; bp, base pairs.

**Supplementary Figure 11. IBD lengths and kinship inference for ancient DNA samples**

(A, C) Scatter plots of IBD lengths estimated by clusIBD and IBIS, TRUFFLE, and IBDseq. Dotted lines represent a line with a slope of 1, i.e., y=x, and points below the line indicate a larger IBD lengths by clusIBD than by IBIS (circle), TRUFFLE (triangle), and IBDseq (square). (B, D) Match matrices of inferred relationships by Fowler et al. and by clusIBD. Different parameters were used, with the default setting for parts A and B, and 300 SNPs per window for parts C and D. IBD, identity by descent; UN, unrelated individuals; Mb, megabases.

**Supplementary Figure 12. Comparison of the running time of clusIBD, IBIS, TRUFFLE, and IBDseq**

All the analyses are performed on an Intel(R) Xeon(R) CPU E5-2650 v4 @ 2.20GHz processor using five cores.

**Supplementary Figure 13. Performance of IBIS when using different acceptable errors**

The figure is generated based on five hundred of simulated 20 Mb segments. Mb, megabases.

**Supplementary Figure 14. Comparison of clusIBD and GEDmatch method for detecting IBD segments of the eighteen artificial poor-quality DNA**

Black dotted lines represent a line with a slope of 1, i.e., y=x, and points below the line indicate a larger IBD lengths (Mb) by clusIBD than by GEDmatch. 150 (default), 200, and 250 SNPs per window were used for parts A, B, and C, respectively. SNPs, single nucleotide polymorphisms; Mb, megabases.

## References

[1] Weir BS, Anderson AD, Hepler AB. Genetic relatedness analysis: Modern data and new challenges. Nat Rev Genet 2006;7:771–80. 10.1038/nrg1960.

[2] Andersen MM, Kampmann M-L, Jepsen AH, Morling N, Eriksen PS, Børsting C, et al. Shotgun DNA sequencing for human identification: Dynamic SNP selection and likelihood ratio calculations accounting for errors. Forensic Sci Int Genet 2025;74:103146. 10.1016/j.fsigen.2024.103146.

[3] Nguyen R, Kapp JD, Sacco S, Myers SP, Green RE. A computational approach for positive genetic identification and relatedness detection from low-coverage shotgun sequencing data. J Hered 2023;114. 10.1093/jhered/esad041.

[4] Voight BF, Pritchard JK. Confounding from cryptic relatedness in case-control association studies. PLoS Genet 2005;1:e32. 10.1371/journal.pgen.0010032.

[5] Liu Y, Bennett EA, Fu Q. Evolving ancient DNA techniques and the future of human history. Cell 2022;185:2632–5. 10.1016/j.cell.2022.06.009.

[6] Petty LE, Phillippi-Falkenstein K, Kubisch HM, Raveendran M, Harris RA, Vallender EJ, et al. Pedigree reconstruction and distant pairwise relatedness estimation from genome sequence data: A demonstration in a population of rhesus macaques (Macaca mulatta). Mol Ecol Resour 2021;21:1333–46. 10.1111/1755-0998.13317.

[7] Stevens EL, Heckenberg G, Roberson EDO, Baugher JD, Downey TJ, Pevsner J. Inference of relationships in population data using identity-by-descent and identity-by-state. PLoS Genet 2011;7:e1002287. 10.1371/journal.pgen.1002287.

[8] Schraiber JG, Akey JM. Methods and models for unravelling human evolutionary history. Nat Rev Genet 2015;16:727–40. 10.1038/nrg4005.

[9] Bieber FR, Brenner CH, Lazer D. Finding criminals through DNA of their relatives. Science 2006;312:1315–6. 10.1126/science.1122655.

[10] Ram N, Guerrini CJ, McGuire AL. Genealogy databases and the future of criminal investigation. Science 2018;360:1078–9. 10.1126/science.aau1083.

[11] Erlich Y, Shor T, Pe’er I, Carmi S. Identity inference of genomic data using long-range familial searches. Science 2018;362:690–4. 10.1126/science.aau4832.

[12] Ge J, Budowle B. Forensic investigation approaches of searching relatives in DNA databases. J Forensic Sci 2021;66:430–43. 10.1111/1556-4029.14615.

[13] Fowler C, Olalde I, Cummings V, Armit I, Büster L, Cuthbert S, et al. A high-resolution picture of kinship practices in an Early Neolithic tomb. Nature 2022;601:584–7. 10.1038/s41586-021-04241-4.

[14] Popli D, Peyrégne S, Peter BM. KIN: a method to infer relatedness from low-coverage ancient DNA. Genome Biol 2023;24:10. 10.1186/s13059-023-02847-7.

[15] Speed D, Balding DJ. Relatedness in the post-genomic era: Is it still useful? Nat Rev Genet 2015;16:33–44. 10.1038/nrg3821.

[16] Zhou Y, Browning SR, Browning BL, Browning BL. IBDkin: Fast estimation of kinship coefficients from identity by descent segments. Bioinformatics 2020;36:4519–20. 10.1093/bioinformatics/btaa569.

[17] Purcell S, Neale B, Todd-Brown K, Thomas L, Ferreira MAR, Bender D, et al. PLINK: A Tool Set for Whole-Genome Association and Population-Based Linkage Analyses. Am J Hum Genet 2007;81:559. 10.1086/519795.

[18] Manichaikul A, Mychaleckyj JC, Rich SS, Daly K, Sale M, Chen W-M. Robust relationship inference in genome-wide association studies. Bioinformatics 2010;26:2867–73. 10.1093/bioinformatics/btq559.

[19] Arciero E, Dogra SA, Malawsky DS, Mezzavilla M, Tsismentzoglou T, Huang QQ, et al. Fine-scale population structure and demographic history of British Pakistanis. Nat Commun 2021;12:7189. 10.1038/s41467-021-27394-2.

[20] Barry A, McNulty MT, Jia X, Gupta Y, Debiec H, Luo Y, et al. Multi-population genome-wide association study implicates immune and non-immune factors in pediatric steroid-sensitive nephrotic syndrome. Nat Commun 2023;14:2481. 10.1038/s41467-023-37985-w.

[21] Manichaikul A, Palmas W, Rodriguez CJ, Peralta CA, Divers J, Guo X, et al. Population structure of hispanics in the United States: The multi-Ethnic study of Atherosclerosis. PLoS Genet 2012;8:e1002640. 10.1371/journal.pgen.1002640.

[22] Kuhn JMM, Jakobsson M, Günther T. Estimating genetic kin relationships in prehistoric populations. PLoS One 2018;13:e0195491. 10.1371/journal.pone.0195491.

[23] Ramstetter MD, Dyer TD, Lehman DM, Curran JE, Duggirala R, Blangero J, et al. Benchmarking relatedness inference methods with genome-wide data from thousands of relatives. Genetics 2017;207:75–82. 10.1534/genetics.117.1122.

[24] Conomos MP, Reiner AP, Weir BS, Thornton TA. Model-free Estimation of Recent Genetic Relatedness. Am J Hum Genet 2016;98:127–48. 10.1016/j.ajhg.2015.11.022.

[25] Browning SR, Browning BL. Identity by Descent Between Distant Relatives: Detection and Applications. Annu Rev Genet 2012;46:617–33. 10.1146/annurev-genet-110711-155534.

[26] Zhou Y, Browning SR, Browning BL. A Fast and Simple Method for Detecting Identity-by-Descent Segments in Large-Scale Data. Am J Hum Genet 2020;106:426–37. 10.1016/j.ajhg.2020.02.010.

[27] Freyman WA, Mcmanus KF, Shringarpure SS, Jewett EM, Bryc K, Auton A. Fast and Robust Identity-by-Descent Inference with the Templated Positional Burrows-Wheeler Transform. Mol Biol Evol 2021;38:2131–51. 10.1093/molbev/msaa328.

[28] Turner SD, Nagraj VP, Scholz M, Jessa S, Acevedo C, Ge J, et al. Evaluating the Impact of Dropout and Genotyping Error on SNP-Based Kinship Analysis With Forensic Samples. Front Genet 2022;13:882268. 10.3389/fgene.2022.882268.

[29] Seidman DN, Shenoy SA, Kim M, Babu R, Woods IG, Dyer TD, et al. Rapid, Phase-free Detection of Long Identity-by-Descent Segments Enables Effective Relationship Classification. Am J Hum Genet 2020;106:453–66. 10.1016/j.ajhg.2020.02.012.

[30] Browning BL, Browning SR. Detecting identity by descent and estimating genotype error rates in sequence data. Am J Hum Genet 2013;93:840–51. 10.1016/j.ajhg.2013.09.014.

[31] Dimitromanolakis A, Paterson AD, Sun L. Fast and Accurate Shared Segment Detection and Relatedness Estimation in Un-phased Genetic Data via TRUFFLE. Am J Hum Genet 2019;105:78–88. 10.1016/j.ajhg.2019.05.007.

[32] Ringbauer H, Huang Y, Akbari A, Mallick S, Patterson N, Reich D. ancIBD - Screening for identity by descent segments in human ancient DNA. BioRxiv Prepr 2023. 10.1101/2023.03.08.531671.

[33] Schmidt DA, Campbell NR, Govindarajulu P, Larsen KW, Russello MA. Genotyping-in-Thousands by sequencing (GT-seq) panel development and application to minimally invasive DNA samples to support studies in molecular ecology. Mol Ecol Resour 2020;20:114–24. 10.1111/1755-0998.13090.

[34] Perry GH, Marioni JC, Melsted P, Gilad Y. Genomic-scale capture and sequencing of endogenous DNA from feces. Mol Ecol 2010;19:5332–44. 10.1111/j.1365-294X.2010.04888.x.

[35] Carpenter ML, Buenrostro JD, Valdiosera C, Schroeder H, Allentoft ME, Sikora M, et al. Pulling out the 1%: Whole-Genome capture for the targeted enrichment of ancient dna sequencing libraries. Am J Hum Genet 2013;93:852–64. 10.1016/j.ajhg.2013.10.002.

[36] Ozga AT, Webster TH, Gilby IC, Wilson MA, Nockerts RS, Wilson ML, et al. Urine as a high-quality source of host genomic DNA from wild populations. Mol Ecol Resour 2021;21:170–82. 10.1111/1755-0998.13260.

[37] Davawala A, Stock A, Spiden M, Daniel R, McBain J, Hartman D. Forensic genetic genealogy using microarrays for the identification of human remains: The need for good quality samples – A pilot study. Forensic Sci Int 2022;334:111242. 10.1016/j.forsciint.2022.111242.

[38] Kaiser J. We will find you: DNA search used to nab Golden State Killer can home in on about 60% of white Americans. Science 2018. 10.1126/science.aav7021.

[39] Caballero M, Seidman DN, Qiao Y, Sannerud J, Dyer TD, Lehman DM, et al. Crossover interference and sex-specific genetic maps shape identical by descent sharing in close relatives. PLoS Genet 2019;15:e1007979. 10.1371/journal.pgen.1007979.

[40] Bherer C, Campbell CL, Auton A. Refined genetic maps reveal sexual dimorphism in human meiotic recombination at multiple scales. Nat Commun 2017;8:14994. 10.1038/ncomms14994.

[41] Li H, Handsaker B, Wysoker A, Fennell T, Ruan J, Homer N, et al. The Sequence Alignment/Map format and SAMtools. Bioinformatics 2009;25:2078–9. 10.1093/bioinformatics/btp352.

[42] Thompson EA. Identity by descent: Variation in meiosis, across genomes, and in populations. Genetics 2013;194:301–26. 10.1534/genetics.112.148825.

[43] Tang K, Naseri A, Wei Y, Zhang S, Zhi D. Open-source benchmarking of IBD segment detection methods for biobank-scale cohorts. Gigascience 2022;11:giac129. 10.1093/gigascience/giac111.

[44] Henn BM, Hon L, Macpherson JM, Eriksson N, Saxonov S, Pe’er I, et al. Cryptic distant relatives are common in both isolated and cosmopolitan genetic samples. PLoS One 2012;7. 10.1371/journal.pone.0034267.

[45] Browning SR, Browning BL. Probabilistic Estimation of Identity by Descent Segment Endpoints and Detection of Recent Selection. Am J Hum Genet 2020;107:895–910. 10.1016/j.ajhg.2020.09.010.

[46] Browning SR, Browning BL. Haplotype phasing: Existing methods and new developments. Nat Rev Genet 2011;12:703–14. 10.1038/nrg3054.

[47] Naseri A, Liu X, Tang K, Zhang S, Zhi D. RaPID: ultra-fast, powerful, and accurate detection of segments identical by descent (IBD) in biobank-scale cohorts. Genome Biol 2019;20:143. 10.1186/s13059-019-1754-8.

[48] Browning SR, Browning BL. Biobank-scale inference of multi-individual identity by descent and gene conversion. Am J Hum Genet 2024;111:691–700. 10.1016/j.ajhg.2024.02.015.

[49] Shemirani R, Belbin GM, Avery CL, Kenny EE, Gignoux CR, Ambite JL. Rapid detection of identity-by-descent tracts for mega-scale datasets. Nat Commun 2021;12:3546. 10.1038/s41467-021-22910-w.

[50] Nait Saada J, Kalantzis G, Shyr D, Cooper F, Robinson M, Gusev A, et al. Identity-by-descent detection across 487,409 British samples reveals fine scale population structure and ultra-rare variant associations. Nat Commun 2020;11:6130. 10.1038/s41467-020-19588-x.

[51] Prüfer K, Stenzel U, Hofreiter M, Pääbo S, Kelso J, Green RE. Computational challenges in the analysis of ancient DNA. Genome Biol 2010;11:R47. 10.1186/gb-2010-11-5-r47.

[52] Fernandes DM, Cheronet O, Gelabert P, Pinhasi R. TKGWV2: an ancient DNA relatedness pipeline for ultra-low coverage whole genome shotgun data. Sci Rep 2021;11:21262. 10.1038/s41598-021-00581-3.

[53] Kennett DJ, Plog S, George RJ, Culleton BJ, Watson AS, Skoglund P, et al. Archaeogenomic evidence reveals prehistoric matrilineal dynasty. Nat Commun 2017;8:14115. 10.1038/ncomms14115.

[54] Hauser SS, Galla SJ, Putnam AS, Steeves TE, Latch EK. Comparing genome-based estimates of relatedness for use in pedigree-based conservation management. Mol Ecol Resour 2022;22:2546–58. 10.1111/1755-0998.13630.

[55] Marsh WA, Brace S, Barnes I. Inferring biological kinship in ancient datasets: comparing the response of ancient DNA-specific software packages to low coverage data. BMC Genomics 2023;24:111. 10.1186/s12864-023-09198-4.

[56] Williams CM, Scelza BA, Daya M, Lange EM, Gignoux CR, Henn BM. A rapid, accurate approach to inferring pedigrees in endogamous populations. BioRxiv Prepr 2020. 10.1101/2020.02.25.965376.

[57] Morimoto C, Manabe S, Fujimoto S, Hamano Y, Tamaki K. Discrimination of relationships with the same degree of kinship using chromosomal sharing patterns estimated from high-density SNPs. Forensic Sci Int Genet 2018;33:10–6. 10.1016/j.fsigen.2017.11.010.

[58] Qiao Y, Sannerud JG, Basu-Roy S, Hayward C, Williams AL. Distinguishing pedigree relationships via multi-way identity by descent sharing and sex-specific genetic maps. Am J Hum Genet 2021;108:68–83. 10.1016/j.ajhg.2020.12.004.

[59] Han E, Carbonetto P, Curtis RE, Wang Y, Granka JM, Byrnes J, et al. Clustering of 770,000 genomes reveals post-colonial population structure of North America. Nat Commun 2017;8:14238. 10.1038/ncomms14238.

[60] Campbell CD, Chong JX, Malig M, Ko A, Dumont BL, Han L, et al. Estimating the human mutation rate using autozygosity in a founder population. Nat Genet 2012;44:1277–81. 10.1038/ng.2418.

[61] Zhou Y, Browning BL, Browning SR. Population-Specific Recombination Maps from Segments of Identity by Descent. Am J Hum Genet 2020;107:137–48. 10.1016/j.ajhg.2020.05.016.

[62] Belbin GM, Odgis J, Sorokin EP, Yee MC, Kohli S, Glicksberg BS, et al. Genetic identification of a common collagen disease in puerto ricans via identity-by-descent mapping in a health system. Elife 2017;6:e25060. 10.7554/eLife.25060.

[63] Gusev A, Kenny EE, Lowe JK, Salit J, Saxena R, Kathiresan S, et al. DASH: A method for identical-by-descent haplotype mapping uncovers association with recent variation. Am J Hum Genet 2011;88:706–17. 10.1016/j.ajhg.2011.04.023.

[64] Browning SR, Thompson EA. Detecting rare variant associations by identity-by-descent mapping in case-control studies. Genetics 2012;190:1521–31. 10.1534/genetics.111.136937.

[65] Temple SD, Waples RK, Browning SR. Modeling recent positive selection using identity-by-descent segments. Am J Hum Genet 2024;111:2510–29. 10.1016/j.ajhg.2024.08.023.

[66] Cai R, Browning BL, Browning SR. Identity-by-descent-based estimation of the X chromosome effective population size with application to sex-specific demographic history. G3 Genes, Genomes, Genet 2023;13:jkad165. 10.1093/g3journal/jkad165.

