## Supplementary figures and images for "clusIBD: Robust Detection of Identity-by-descent (IBD) Segments Using Unphased Genetic Data from Poor-quality DNA Samples"

### Supplementary Figure 1. distribution of Roph.jpeg

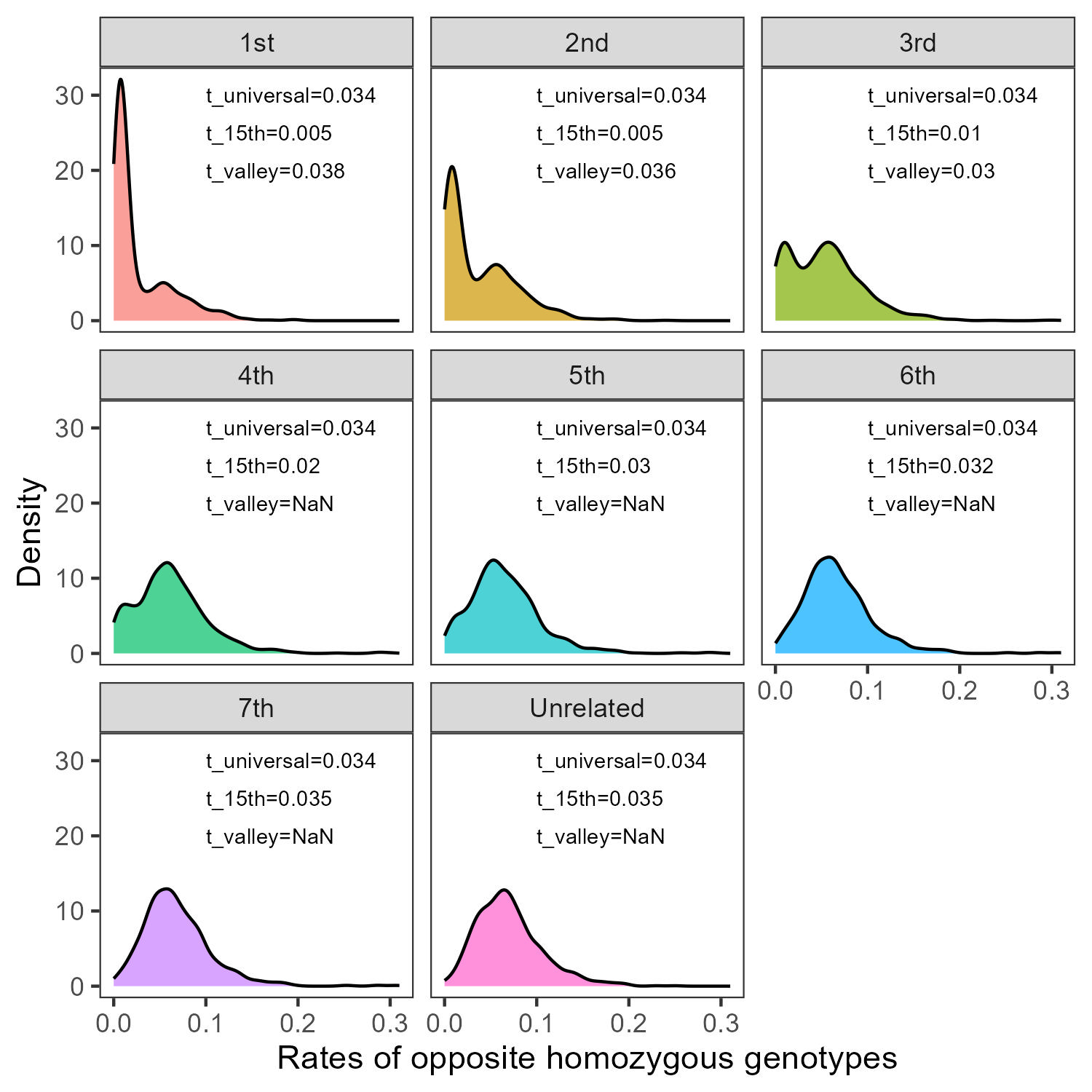

### Supplementary Figure 2. The pedigree of studied family.jpg

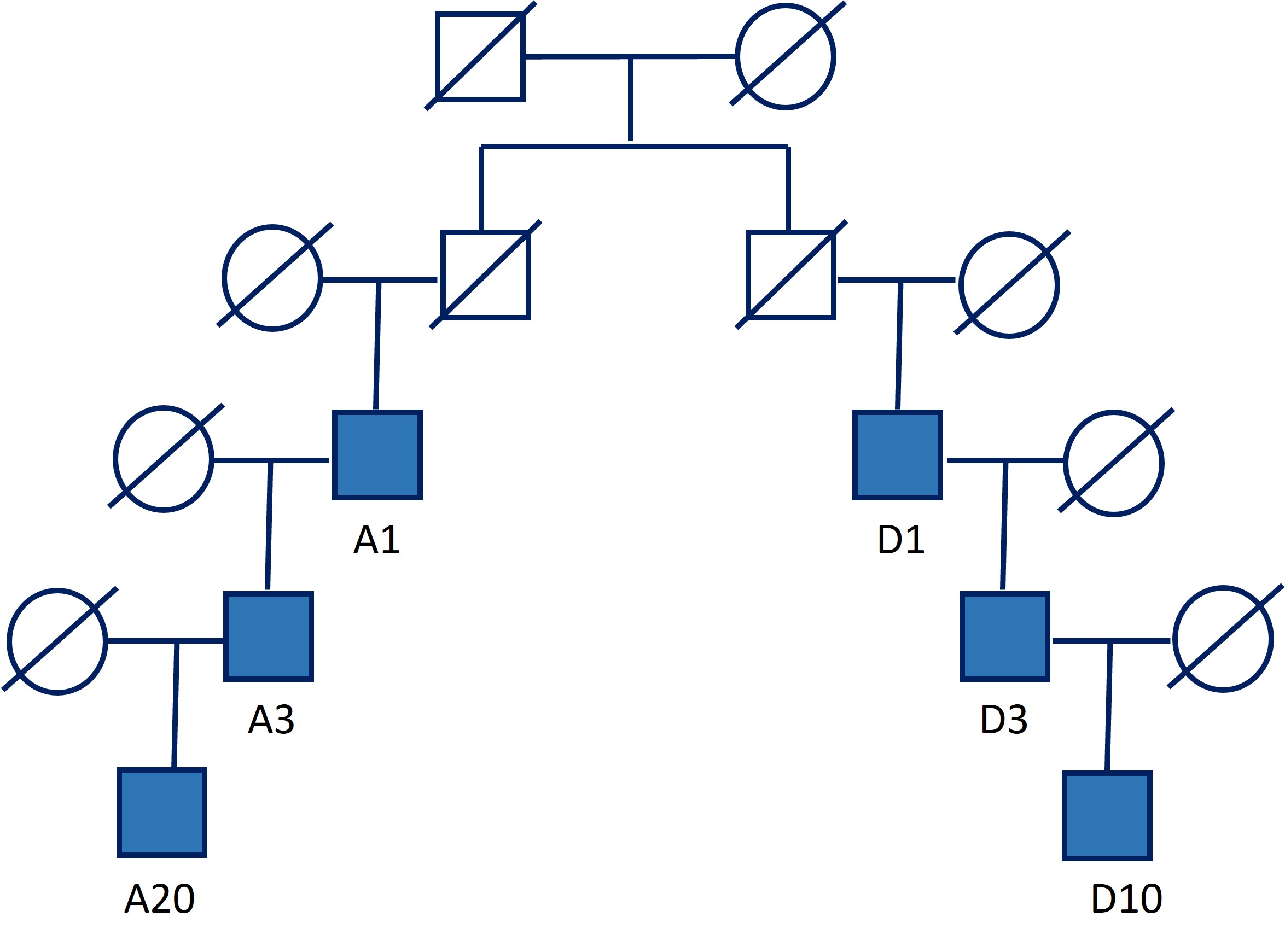

### Supplementary Figure 3 sensitivity study.jpeg

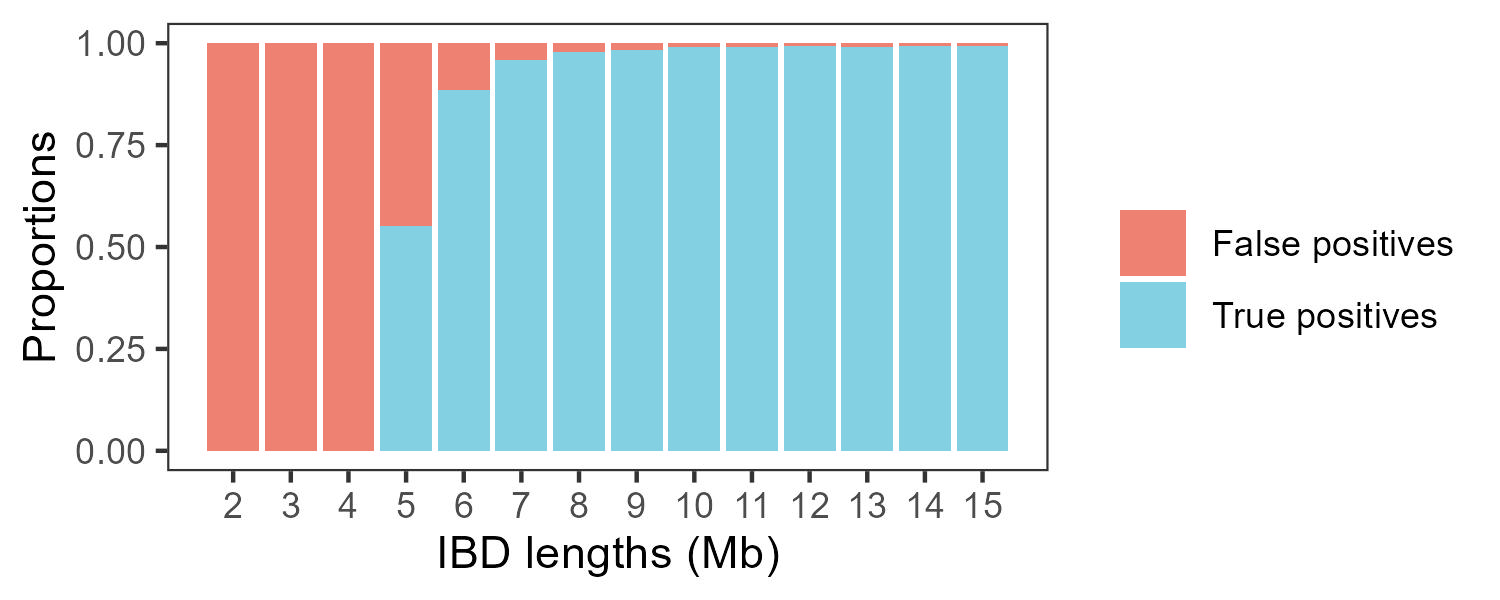

### Supplementary Figure 4. The number of SNPs per window.jpeg

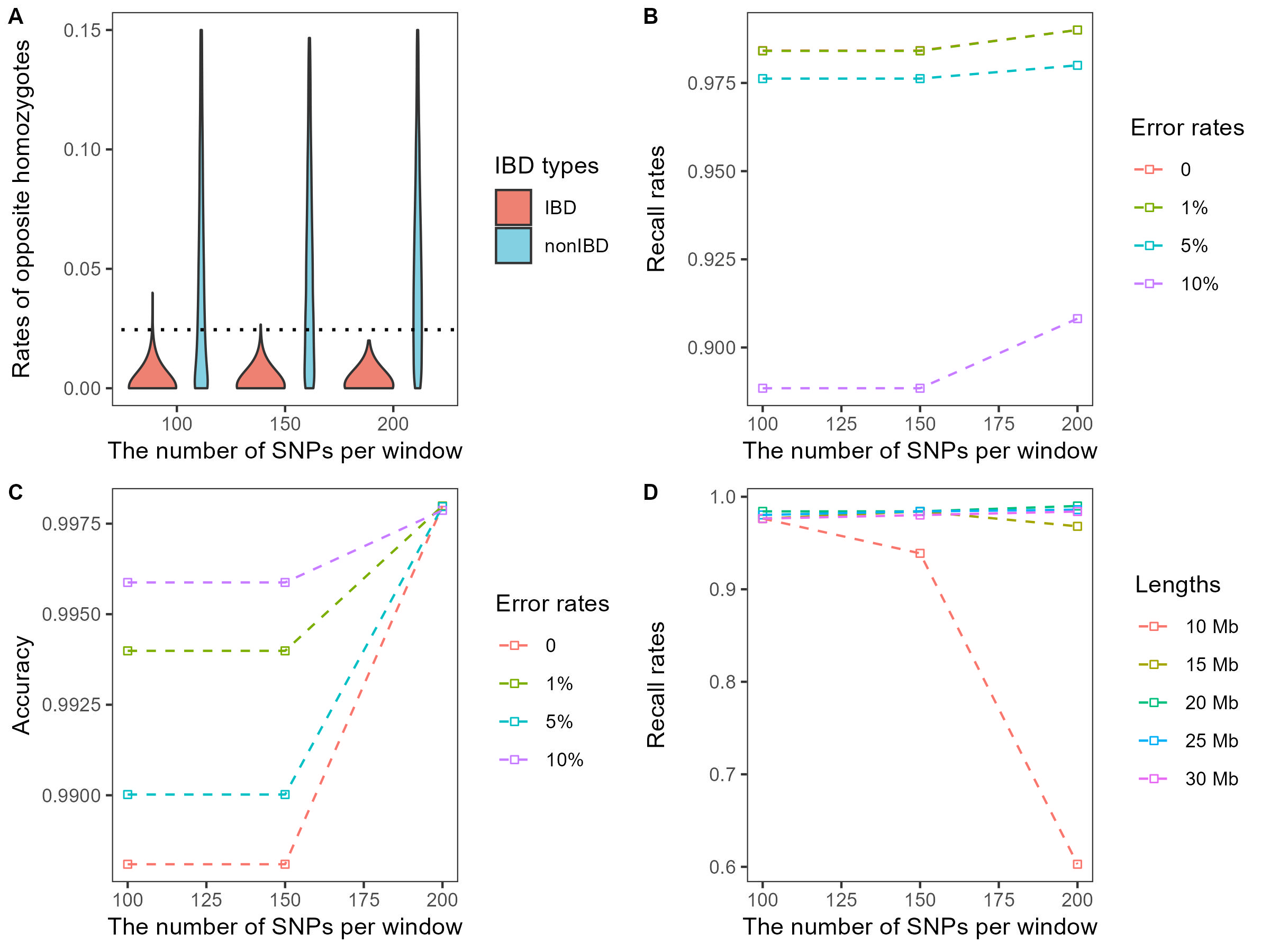

### Supplementary Figure 5. perforamce zoom 0.75-1.jpeg

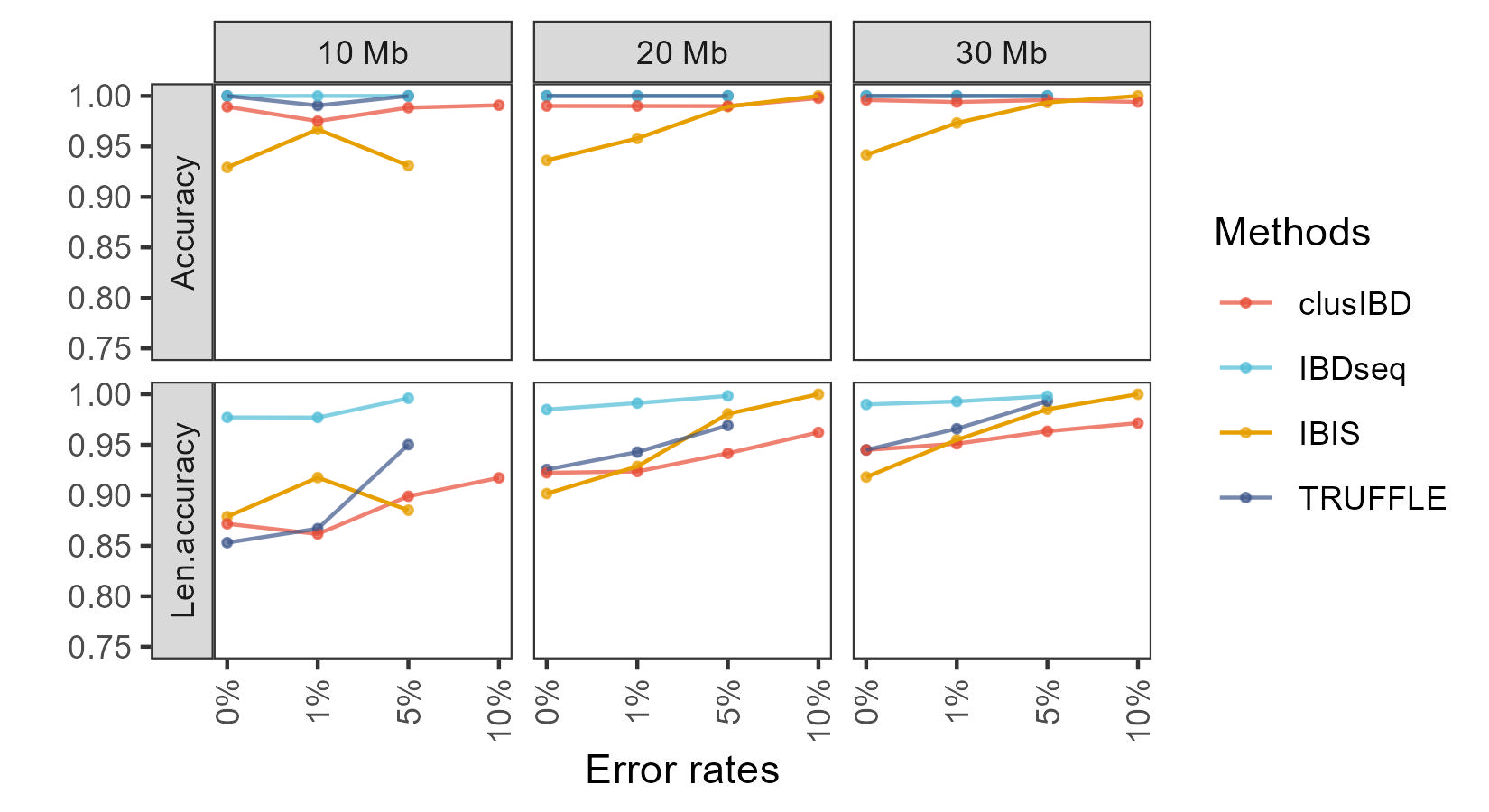

### Supplementary Figure 6. edges of clusIBD.jpeg

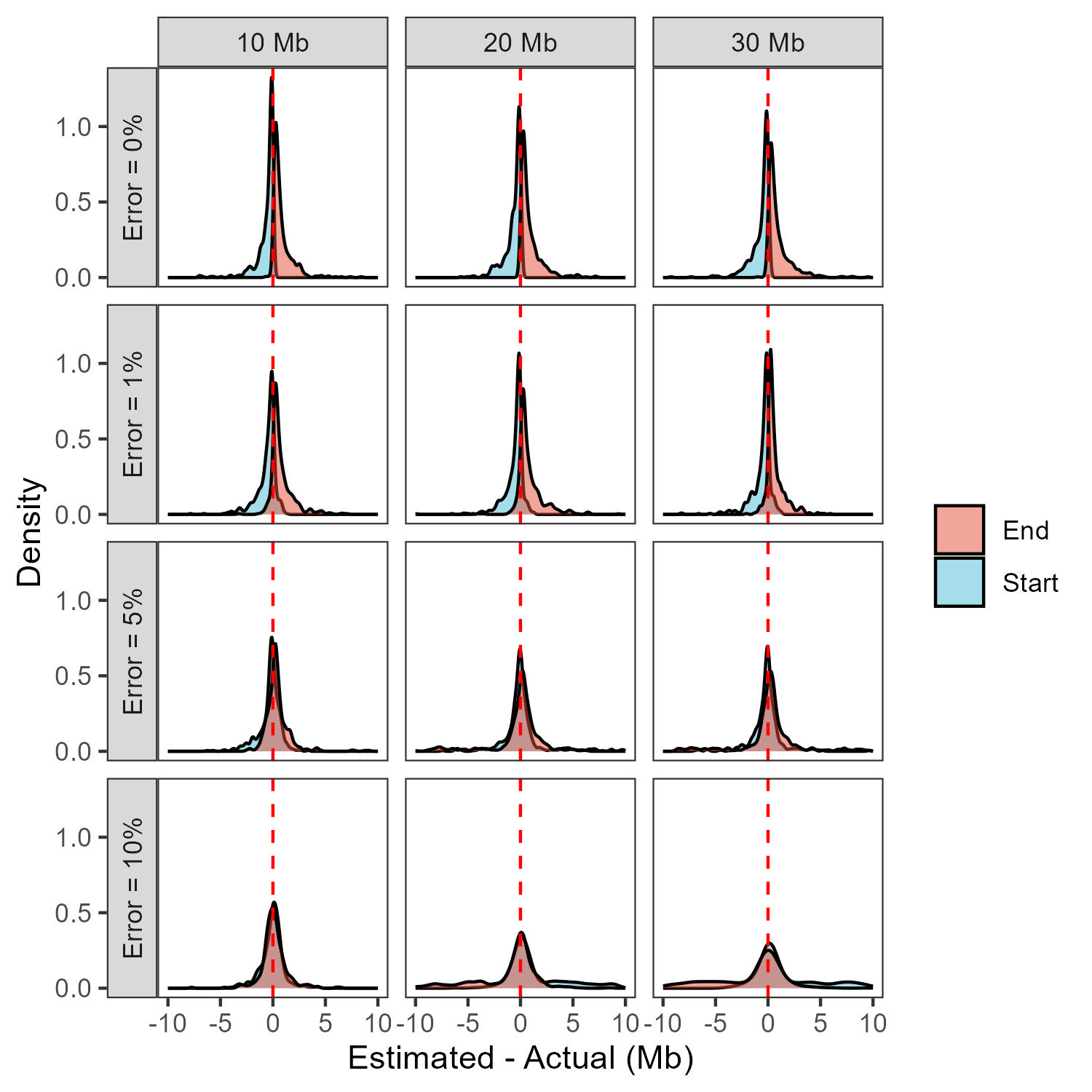

### Supplementary Figure 7. performance on relationships.jpeg

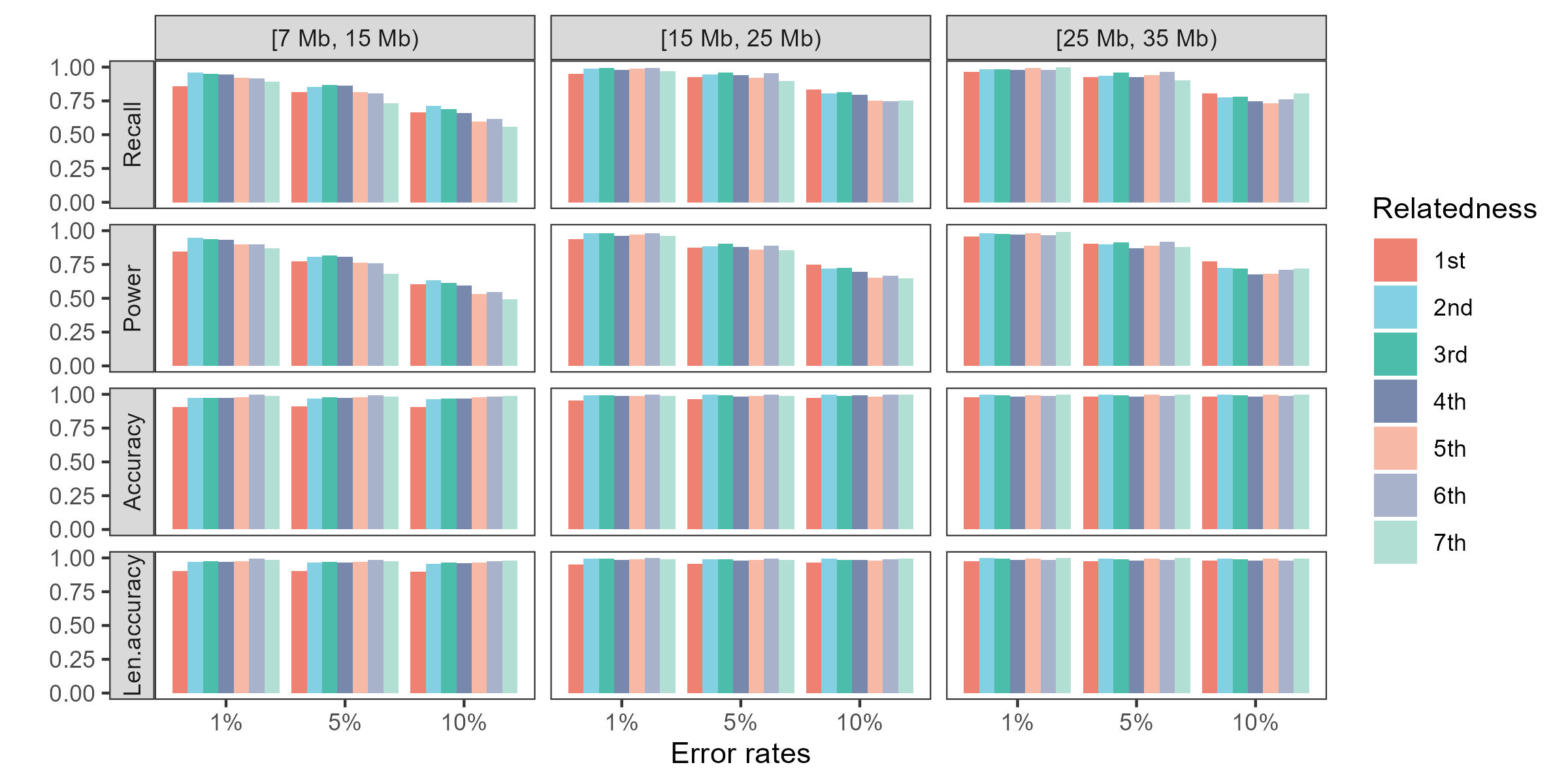

### Supplementary Figure 8. performance on populaitions.jpeg

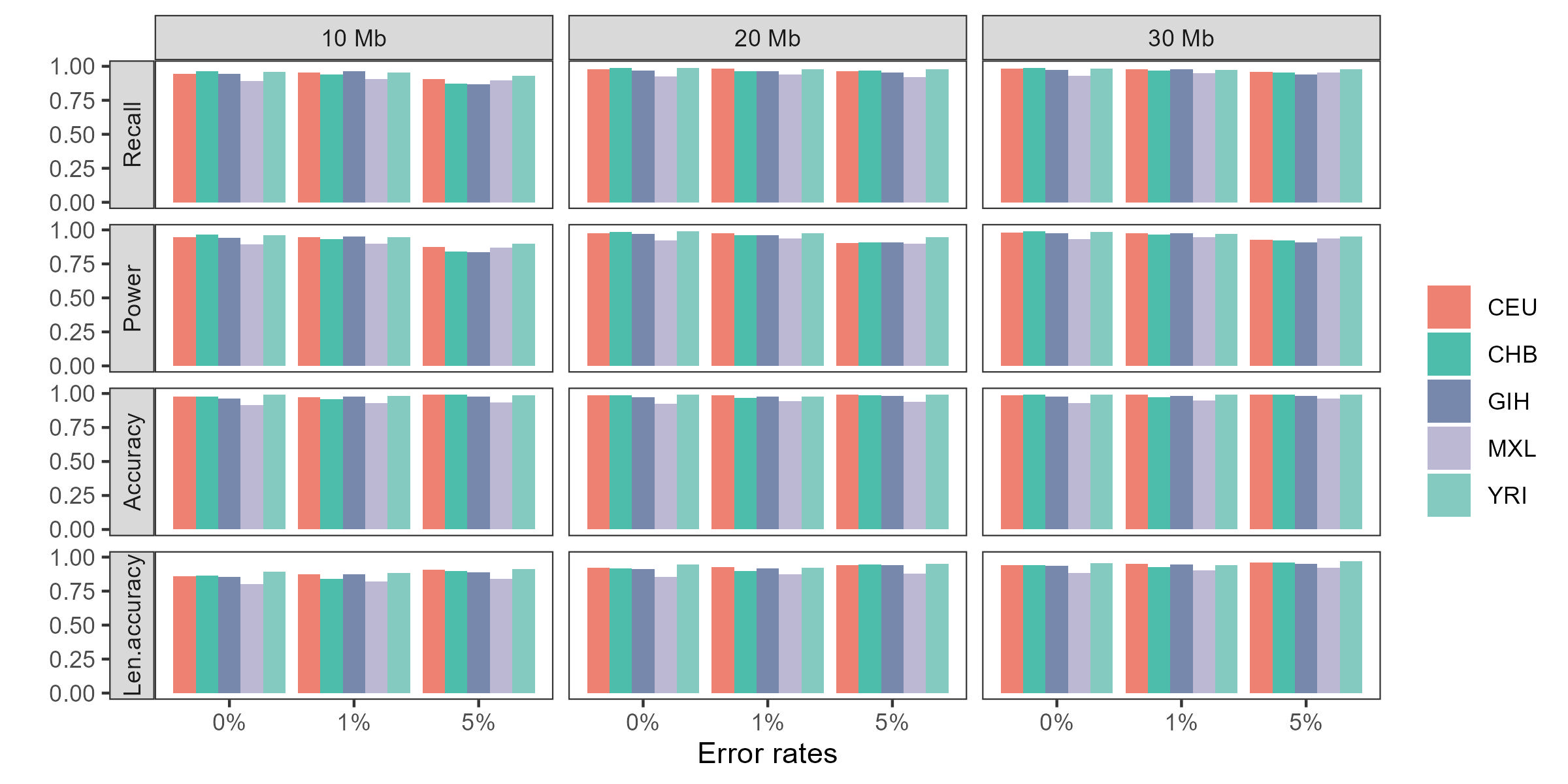

### Supplementary Figure 9. kinship coefficients.jpeg

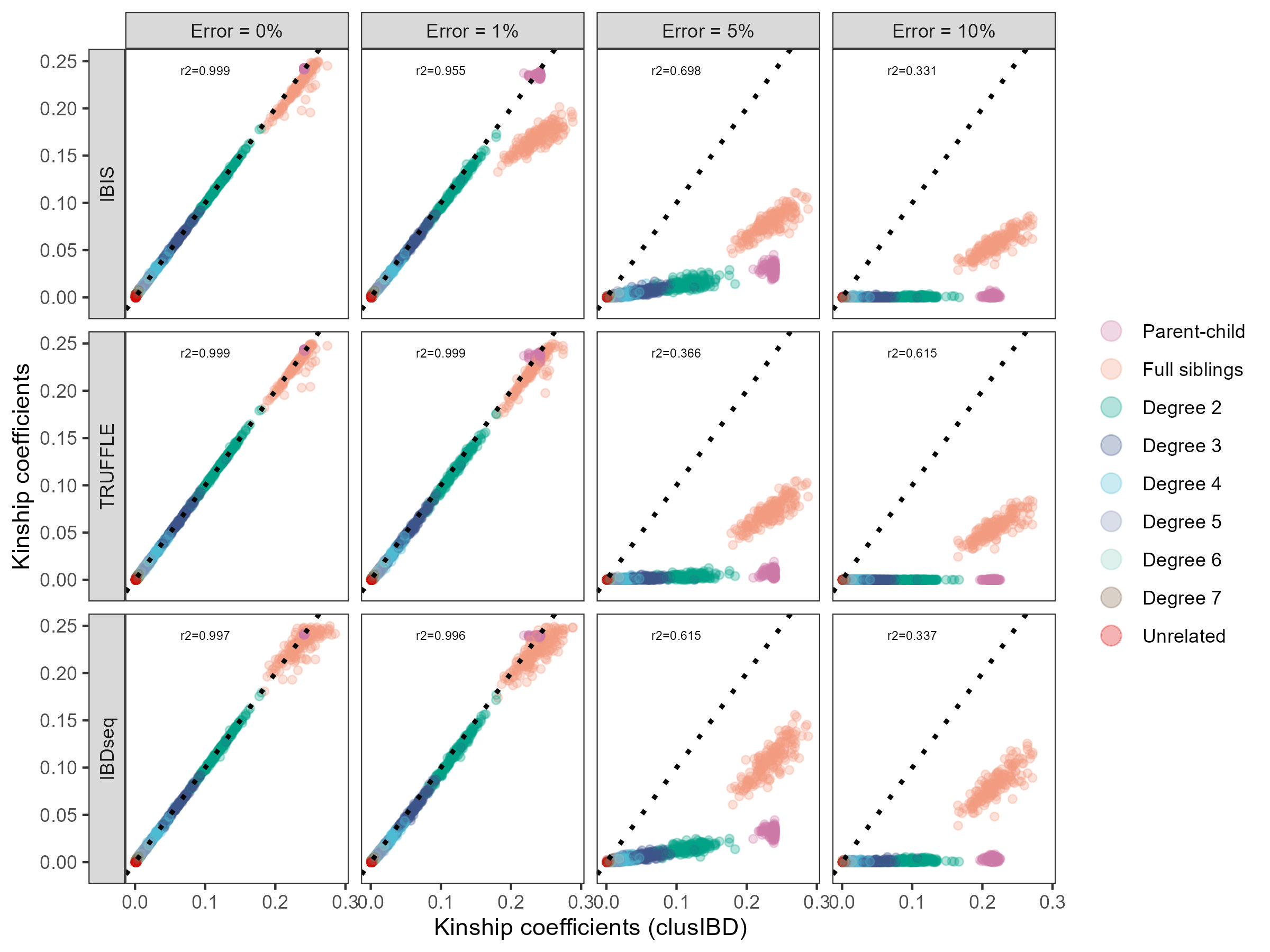

### Supplementary Figure 10. errors of poor quality DNA.jpeg

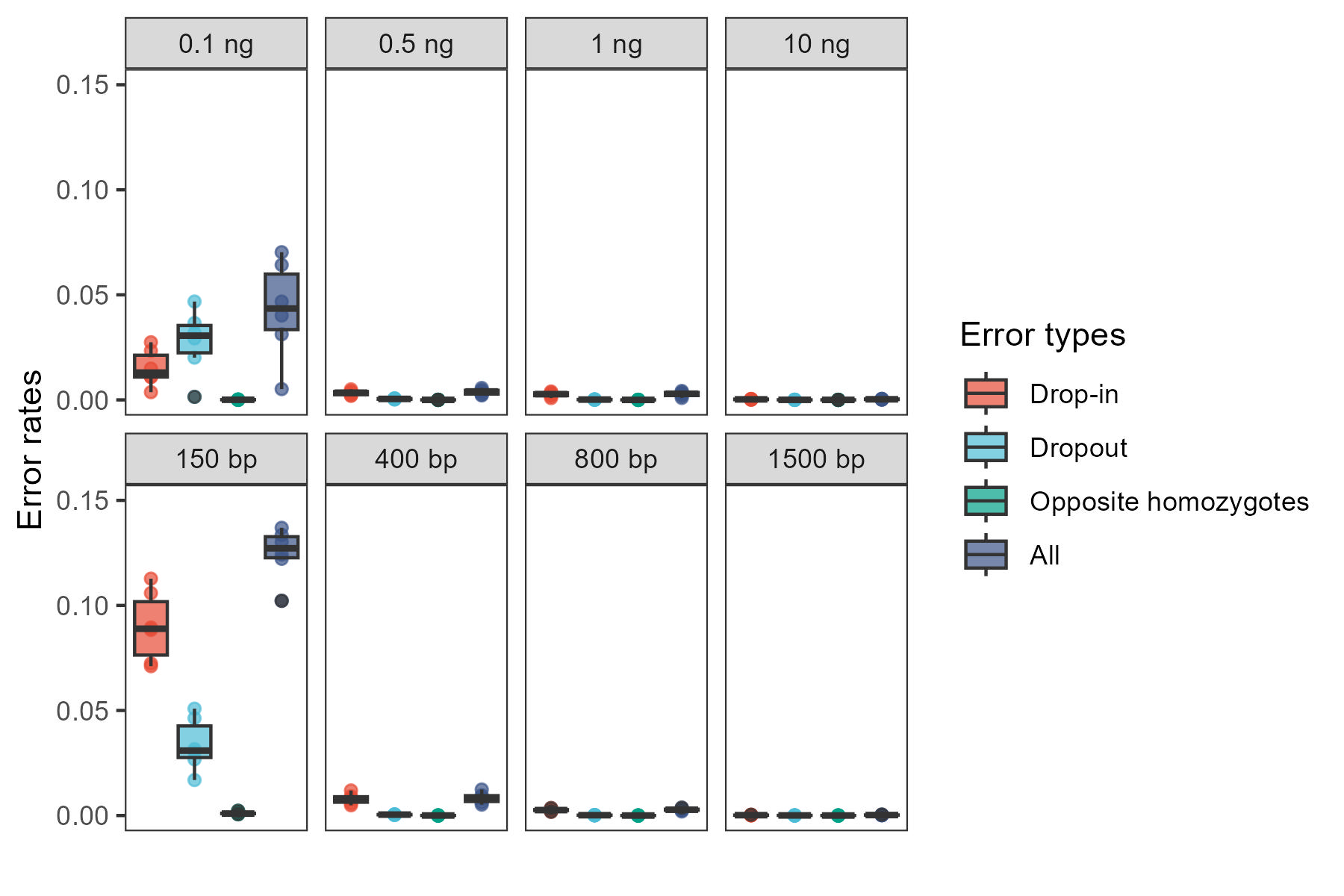

### Supplementary Figure 11. ancientDNA DNA.jpeg

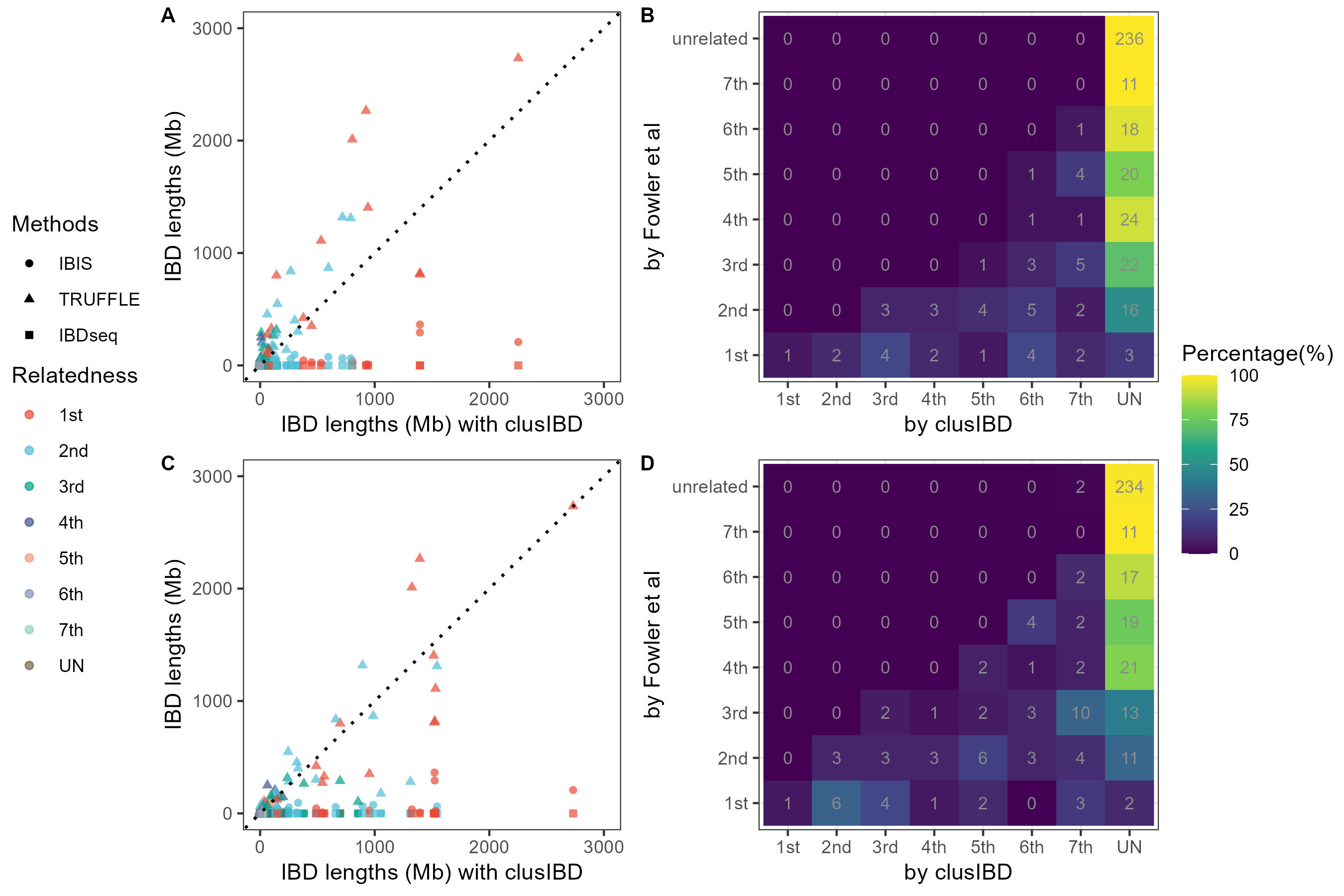

### Supplementary Figure 12. run time.jpeg

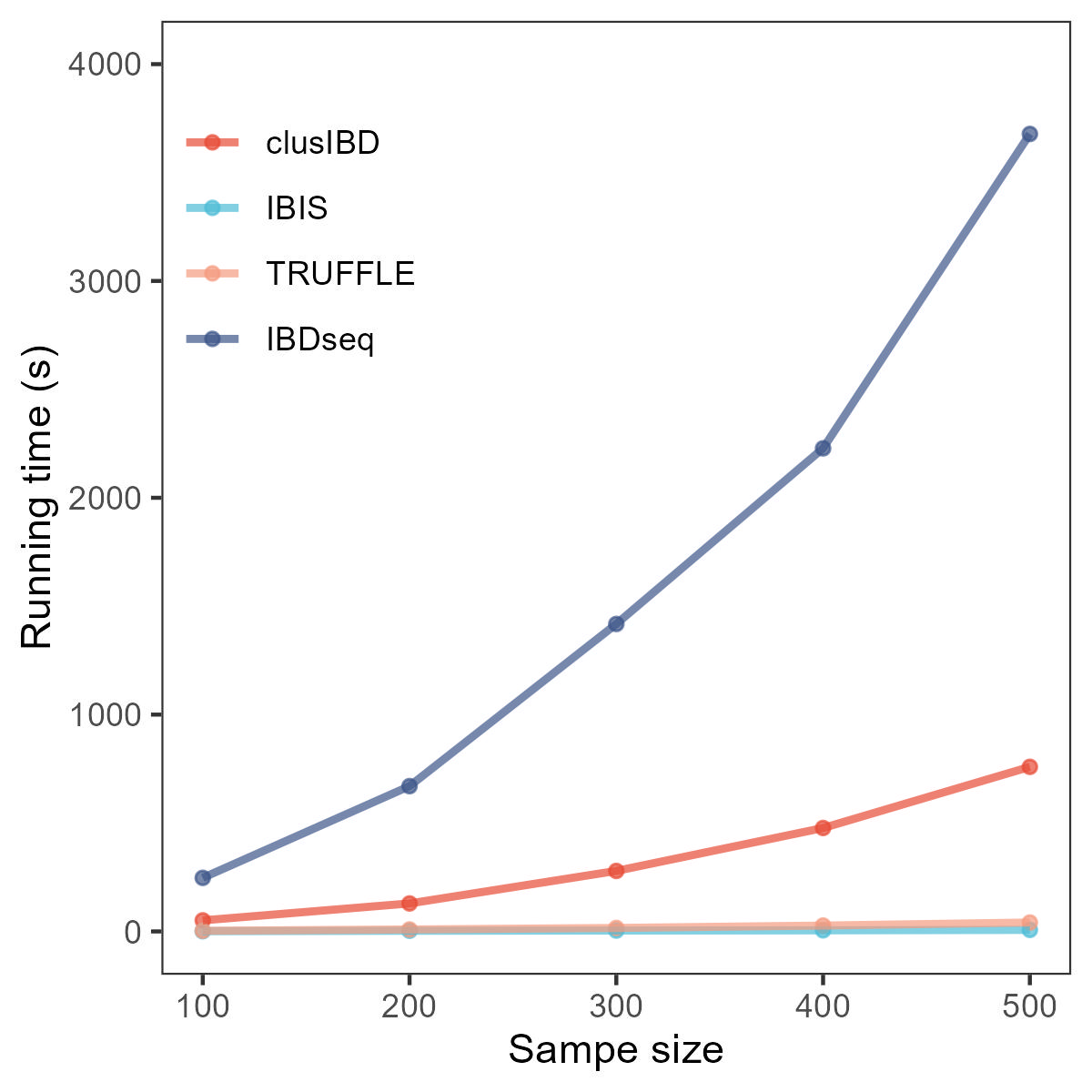

### Supplementary Figure 13. performance with IBIS.jpeg

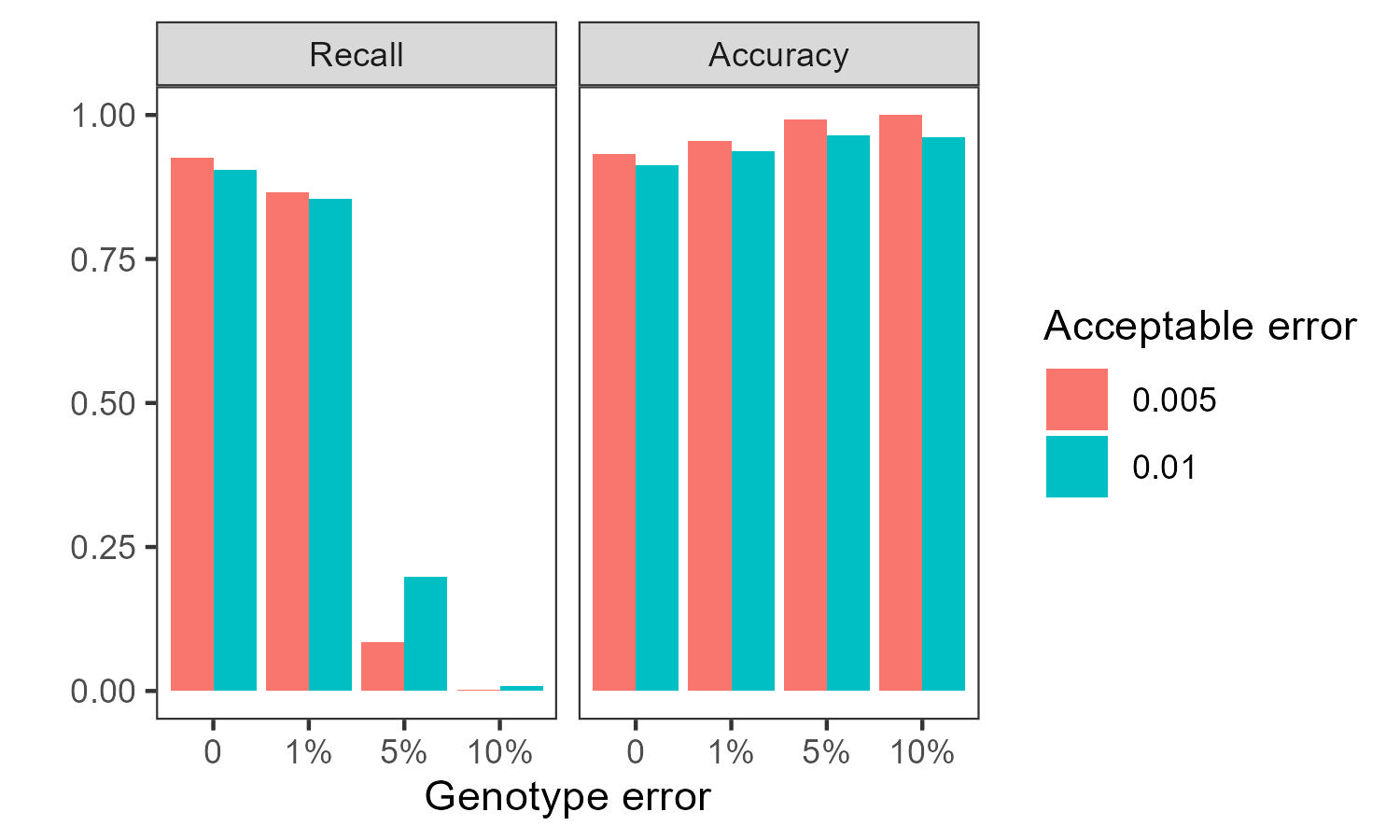

### Supplementary Figure 14. gedmatch and clusIBD.jpeg

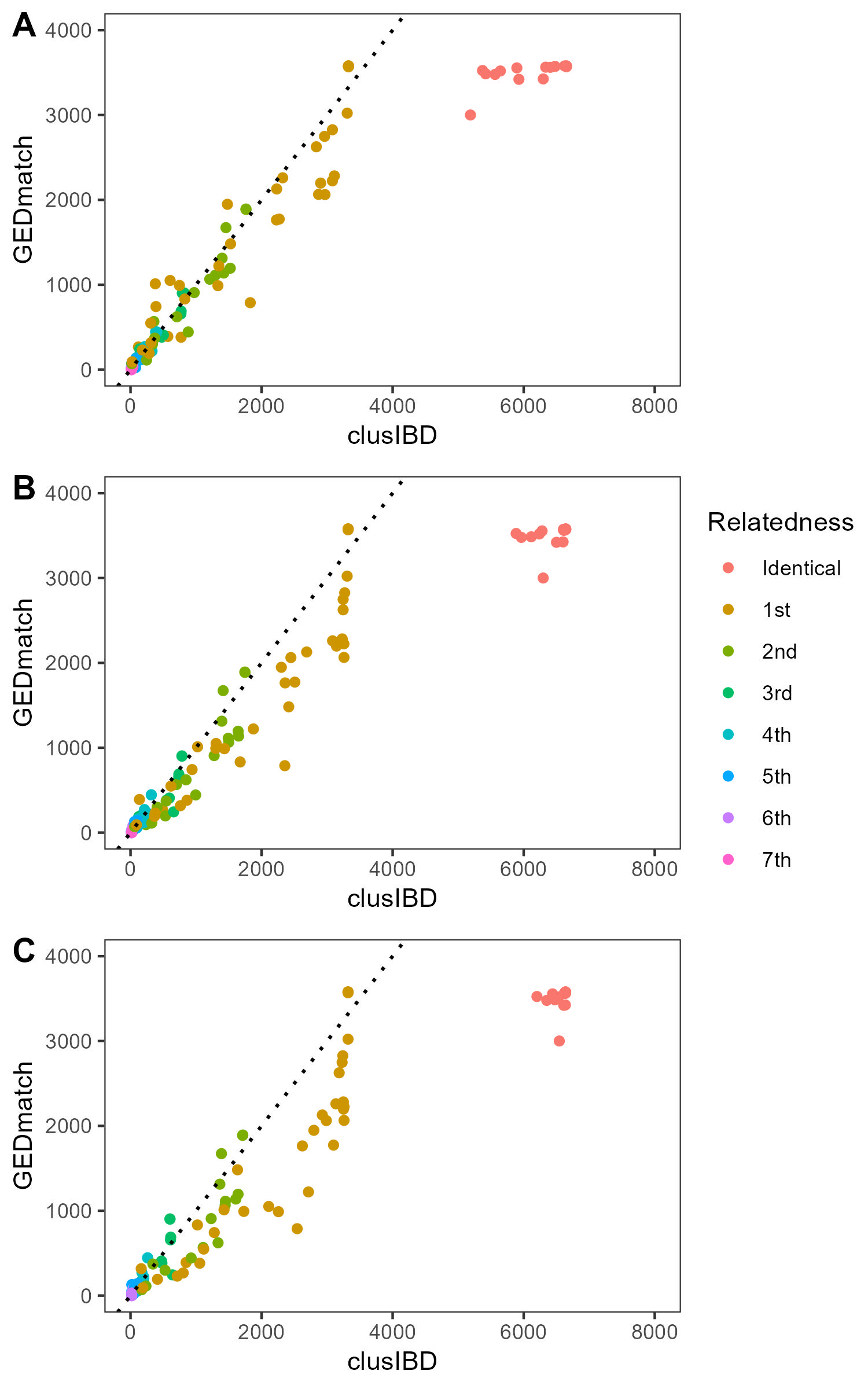
